# Functional interrogation of osteoarthritis-associated regulatory variants and effector genes

**DOI:** 10.64898/2026.11.23.768666

**Authors:** Ana P Piedade, Samantha R Stuppy, Jacqueline Shine, Emma R Klein, Jake E Butler, Antonios Frantzeskos, Justin Engelbert, Sarah J Rice, Stephen Eyre, Jonathan D Rosen, Richard F Loeser, Leela C Biant, Douglas H Phanstiel, Brian O Diekman, Gisela Orozco

## Abstract

**Background:** Genome-wide association studies (GWAS) have identified hundreds of loci associated with osteoarthritis (OA); however, the regulatory variants and effector genes underlying most associations remain unresolved.

**Results:** We used complementary functional genomic approaches in chondrocytes to resolve prioritized OA risk loci. Massively parallel reporter assays (MPRAs) in primary human chondrocytes identified allele-specific regulatory activity at variants across multiple loci, two of which were independently validated using low-throughput reporter assays. In parallel, integration of OA GWAS signals with chromatin accessibility, H3K27ac CUTCTag, and chromatin conformation data prioritised candidate regulatory elements for CRISPR-based perturbation. CRISPR activation and interference supported functional regulatory relationships between OA risk regions and Pappalysin 1 (*PAPPA*) and Sprouty RTK signaling antagonist 4 (*SPRY4*) as target genes. At the 9q33 locus, multiple independent lines of evidence converged on rs1895062, which resides in an accessible region that loops to the *PAPPA* promoter. The OA risk “G” allele at rs1895062 results in higher regulatory activity by MPRA and low throughput reporter assay. Activation of this region by CRISPR increases *PAPPA* gene expression and induces widespread transcriptional changes involving extracellular matrix remodelling and other OA-relevant pathways.

**Conclusion:** Integrating genetic association, genomic and epigenomic data, and functional perturbation in disease-relevant cells can resolve the regulatory mechanisms underlying OA risk loci. Using this multi-faceted approach, we provide convergent functional evidence for rs1895062 as a likely causal variant that controls *PAPPA* gene expression and OA risk.

## Background

The risk for common, complex diseases such as osteoarthritis (OA) arise from the interaction of environmental factors with a large number of genetic variants with modest individual contributions^1,2^. OA is a highly prevalent disease characterised by pain and degradation of articular cartilage and other tissues of the joint^3–5^. Increasingly powered genome-wide association studies (GWAS) have identified over 900 independent genetic associations for OA^6,7^. However, GWAS are only a starting point for deriving mechanistic insight^8,9^. Most disease-associated variants are located within non-coding regions of the genome and are enriched in tissue-specific regulatory elements, which demonstrates the need to evaluate how putative target genes are regulated in disease-relevant cell types^10^. Additionally, a single GWAS signal often encompasses multiple single nucleotide polymorphisms (SNPs) in strong linkage disequilibrium (LD), resulting in highly correlated association signals that cannot be disentangled using only statistical approaches^11^. Given these challenges, developing and testing variant-to-gene hypotheses benefits from the integration of multi-omic datasets in patient-derived tissues and complementary experimental approaches^12,13^.

One functional genomics approach is to integrate genetic and transcriptomic data through expression quantitative trait locus (eQTL) mapping, which links genetic variation to changes in gene expression and thereby identifies putative regulatory variants and disease effector genes. We recently mapped eQTLs in primary human chondrocytes under baseline conditions and in response to stimulation with a fibronectin fragment matrikine that initiates an OA phenotype^14^. Colocalization with OA GWAS signals can further prioritise loci likely to contribute to disease risk through altered gene regulation. However, eQTL analyses provide statistical evidence of regulatory effects rather than direct experimental confirmation of enhancer function.

Massively parallel reporter assays (MPRAs) have successfully identified active enhancer sequences and expression-modulating variants (emVars) across numerous diseases and cellular systems, providing a scalable framework for moving from genetic association to functional validation^15,16^. Despite the power of these advances, OA-associated loci have not been systematically evaluated using MPRA in cells derived from joint tissues. While early candidate regions associated with OA were tested in an osteosarcoma cell line^17^, many regulatory variants exhibit effects that are only detectable in particular cell types or contexts^18–20^. Chondrocytes are a critical cell type when interpreting the genetic risk for OA, as they are the sole cell type in articular cartilage and contribute to homeostasis of the joint^21^. When cultured in high density monolayers for short periods, primary chondrocytes retain key phenotypic and transcriptional characteristics of native cartilage while maintaining the capacity to mount a robust response to OA stimuli^22^. Thus, primary chondrocytes provide a physiologically relevant model for investigating the function of OA-associated variants.

The regulatory relationship between enhancers and promoters is complex and often involves long-range interactions mediated by DNA loops^23^. Previous work by Bittner *et al.* used chromatin conformation mapping in primary OA chondrocytes to investigate the regulatory architecture of OA GWAS loci and identify interactions between putative regulatory elements and candidate target gene promoters^24^. This work generated specific variant-to-gene hypotheses, including the presence of a long-range regulatory relationship at the 9q33 locus. The lead OA GWAS variant rs1321917 is located within an intron of Astrotactin 2 (*ASTN2*), yet the associated risk region interacts with the Pappalysin 1 (*PAPPA*) promoter more than 400 kb away, prioritising it as the likely OA effector gene^14,24^. These findings illustrate how integration of genetic association and chromatin conformation can connect non-coding disease-associated regions to distal candidate effector genes. However, chromatin interactions alone do not establish regulatory activity or demonstrate that a candidate element functionally controls the predicted target gene. Integrating chromatin conformation with disease-relevant epigenomic profiling and experimental perturbation therefore provides complementary approaches for refining candidate regulatory elements and experimentally testing variant-to-gene hypotheses.

Building on these previous genetic and genomic studies, we sought to functionally resolve prioritized OA risk loci using complementary approaches to interrogate regulatory activity and variant-to-gene relationships. We integrated OA GWAS signals with primary chondrocyte eQTLs to prioritize candidate variants for MPRA, followed by low throughput reporter assays to validate selected allele-specific regulatory effects. In parallel, OA GWAS signals were integrated with chromatin accessibility, enhancer activity, and chromatin conformation data to prioritize candidate regulatory elements for CRISPR-based activation and interference. Together, these complementary strategies provide a framework for moving from genetic association toward functional characterization of regulatory variants, effector genes, and molecular mechanisms underlying OA susceptibility.

## Results

We used a combination of genomic and epigenomic datasets to prioritize high-confidence OA risk loci for functional follow-up (Figure 1). The first path prioritized ten loci through colocalization of primary chondrocyte eQTLs and OA GWAS^14^. In parallel, a second path prioritized four variants at three loci through existing chondrocyte Hi-C data^24^ along with newly generated ATAC-seq and H3K27ac CUTCTag data. Each of these three loci was investigated by experimental follow-up, with the most in-depth characterisation performed on loci harbouring the genes *PAPPA* and *SLC44A2* due to prioritization from both bioinformatic pathways (Figure 1).

**Figure 1.**
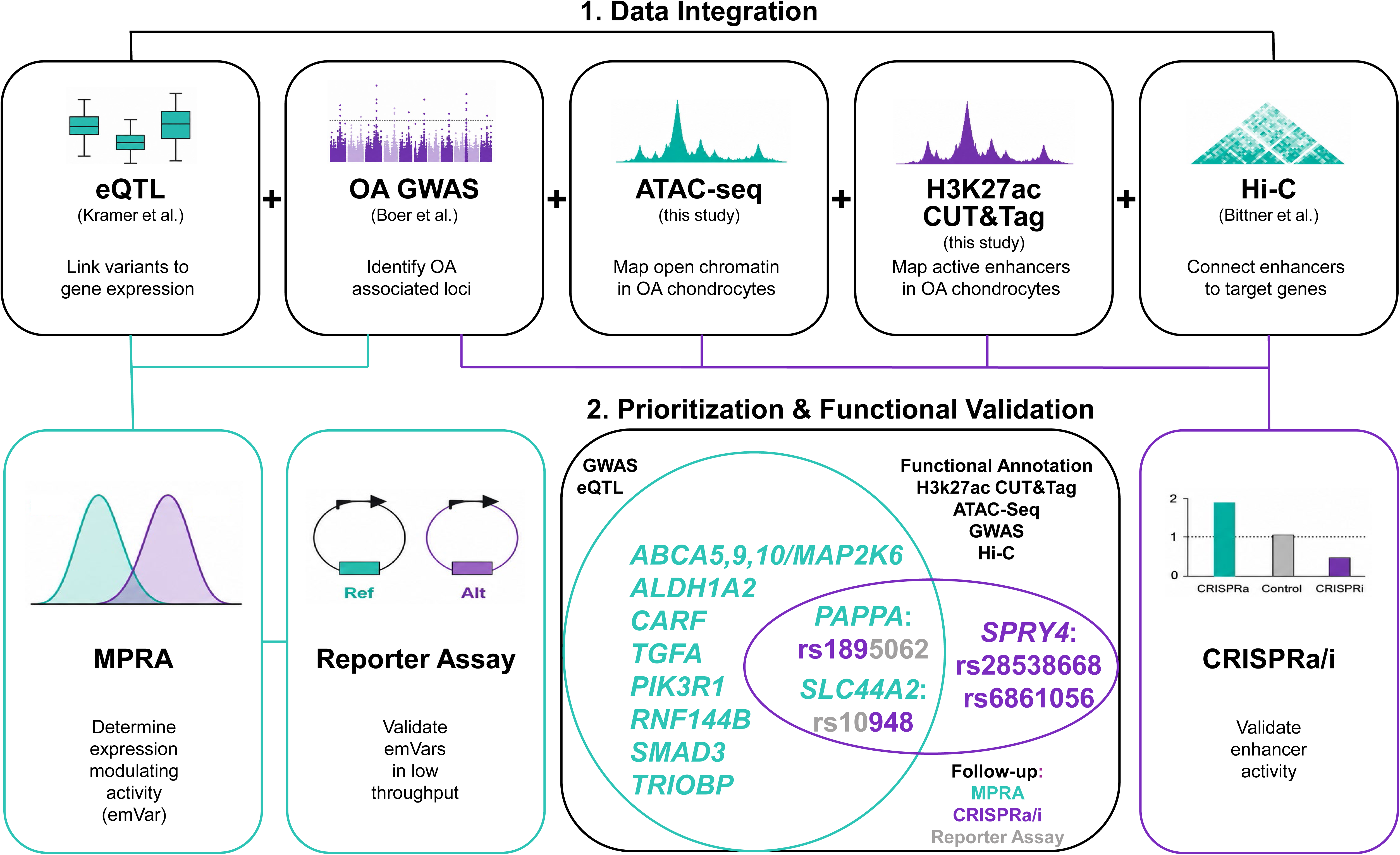
Paths for OA GWAS locus prioritization. OA GWAS signals were integrated with primary chondrocyte eQTLs to prioritize candidate variants for MPRA, with selected allele-specific effects validated by low-throughput reporter assays. In parallel, OA GWAS signals were integrated with chromatin accessibility (ATAC-seq), enhancer activity (H3K27ac CUT&Tag), and chromatin conformation (Hi-C) to prioritize regulator elements for CRISPRa / CRISPRi perturbation.

### Massively parallel reporter assays (MPRAs)

MPRAs are ideally suited to interrogate multiple regions for regulatory activity simultaneously. Restricting the experimental design to OA GWAS loci with significant eQTLs in chondrocytes was selected as an efficient strategy to prioritize loci that are likely to contain emVars. Testing the lead eQTLs plus LD-linked variants at ten prioritized loci provided a moderate MPRA library compatible with use in primary human chondrocytes. The library consisted of 832 test oligos and 300 control oligos (Supplementary Table 1).

We first validated assay performance using three control classes: scrambled negative controls expected to have minimal regulatory activity, previously validated active controls expected to drive reporter expression, and previously identified emVar controls with established allele-specific activity in published MPRA datasets. Negative control sequences exhibited low, tightly centred activity distributions, whereas active controls showed the expected increase in activity (Fig. 2A). OA-associated test sequences spanned a broader range overlapping both control classes, compatible with the expectation that only a subset of tested variants would show detectable activity (Fig. 2A). To assess the concordance of allelic effects detected in primary chondrocytes with MPRA experiments in other cell types, we compared allele-specific emVar activity in a set of 50 variants previously shown to be significant emVars in three cell types^25^. Allelic log₂ fold changes showed modest concordance across datasets, indicating reproducible regulatory effects while also revealing cell type specific activity detectable in primary chondrocytes (Fig. 2B).

**Figure 2.**
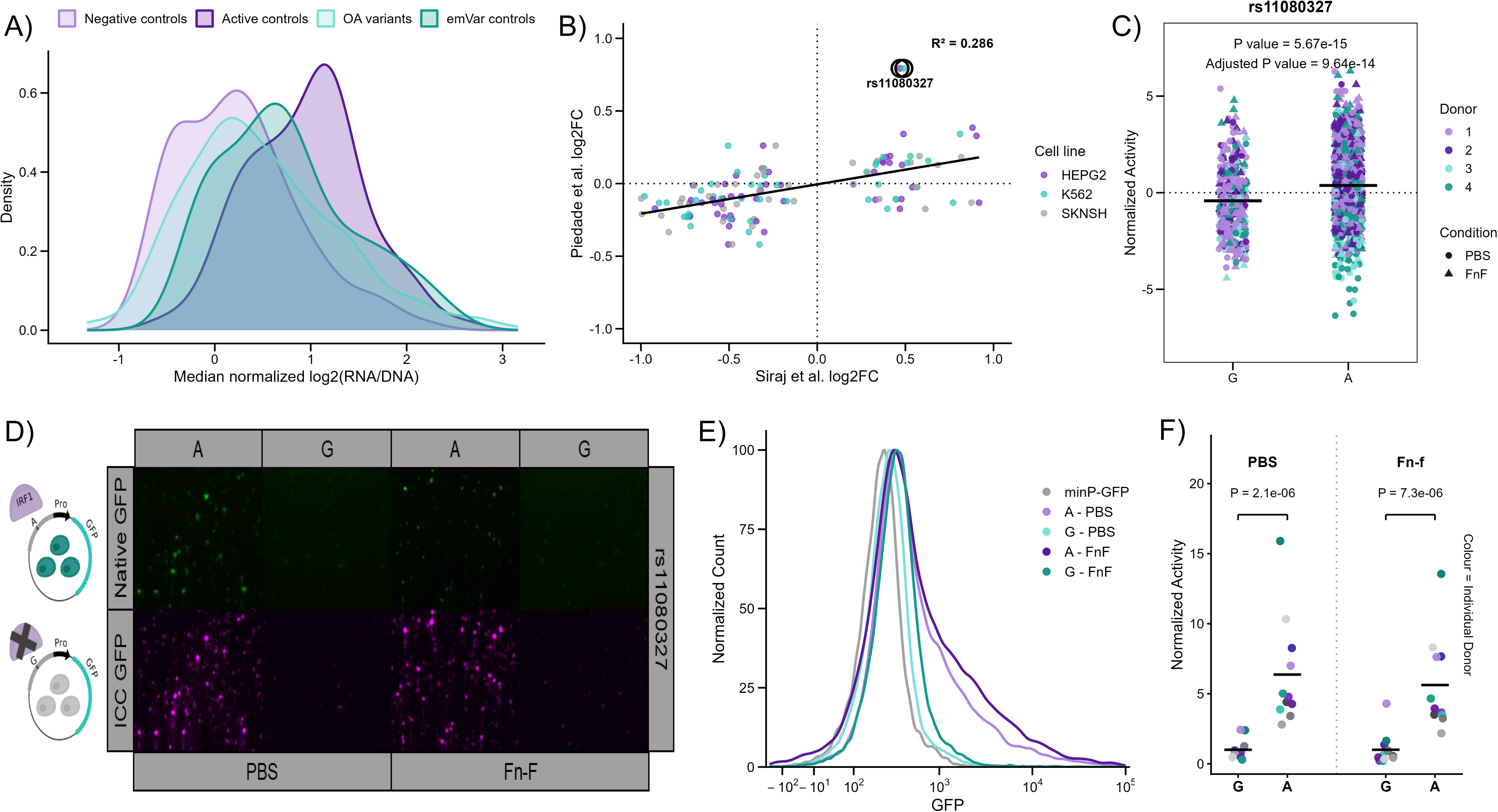
Performance and validation of MPRA in normal primary human chondrocytes. **(A)** Distribution of normalised MPRA activity for negative control sequences, active control sequences, emVar control sequences, and candidate osteoarthritis (OA)–associated variants assayed in primary human chondrocytes (n = 4 donors). **(B)** Comparison of allele-specific MPRA activity estimates obtained in primary human chondrocytes with previously published MPRA datasets from other cellular contexts. Points represent individual variant allele pairs; values correspond to allelic log₂ fold changes. Allelic effects show modest concordance across datasets (Pearson r² = 0.286). **(C)** Allele-specific MPRA activity for rs11080327 at the *SLFN5* locus. Each point represents an individual barcode aggregated across donors; centre line denotes the median. The alternative allele exhibits significantly higher reporter activity than the reference allele (adjusted p = 9.64 x 10^-14^, two-sided t-test following one-pass Grubbs outlier removal). **(D)** Representative immunocytochemistry images of primary human chondrocytes transfected with GFP reporter constructs carrying the reference or alternate allele of rs11080327 under baseline (PBS) and fibronectin fragment (Fn-f) stimulation. **(E)** Representative flow cytometric histograms from a single donor showing GFP reporter activity for rs11080327 constructs. **(F)** Quantification of the fraction of GFP-high cells for rs11080327 reporter constructs across donors and replicates (n = 11). Points represent individual donor measurements. The alternative A allele significantly increased the fraction of GFP-high cells under both PBS and Fn-f conditions (PBS adjusted p = 2.1 x 10^-6^, Fn-f adjusted p = 7.3 x 10^-6^; paired ratio t-test).

Among the control variants tested, rs11080327 at the *SLFN5* locus exhibited a robust allelic effect, with the alternate allele driving significantly higher reporter activity than the reference allele across donors, conditions, and barcodes (Fig. 2C). To validate this MPRA-identified effect, we independently cloned the reference and alternate alleles upstream of the promoter–GFP reporter and assessed reporter activity in primary chondrocytes under baseline (PBS) conditions and after stimulation with a fibronectin fragment (Fn-f), a bioactive matrikine found in OA joints that stimulates an OA-like phenotype in monolayer culture^22^. Immunofluorescence (IF) demonstrated increased GFP signal driven by the alternate “A” allele for rs11080327 relative to the reference “G” allele in both conditions (Fig. 2D). Flow cytometric analysis further confirmed that the alternate allele increased the extent to which cells expressed the reporter gene under both baseline and stimulated conditions in each of 11 independent donors (Fig. 2E–F). Together, these data demonstrate that MPRA reproducibly detects allele-dependent regulatory activity in primary human chondrocytes and that these effects are concordant with independent low-throughput reporter assays.

### MPRA and low throughput follow-up provide support for PAPPA and SLC44A2 variants

Across the ten prioritized loci tested, we identified emVars in regulatory regions linked to 7 eGenes: *CARF, PAPPA, RNF144B, SLC44A2, SMAD3, TGFA, and TRIOBP* (Supplementary Tables 2-4). Several of these eGenes had multiple emVars and the combined-condition analysis identified more emVars than either condition alone. For three tested loci we did not detect any significant emVars: *ALDH1A2, PIK3R1,* and a single eQTL locus associated with the expression of four eGenes (*ABCA5, ABCAS, ABCA10, MAP2KC*). Of the 15 emVars observed, we focused on rs1895062 for *PAPPA* and rs10948 for *SLC44A2* as these putative causal variants were also prioritized by the parallel path of integrated genomic and epigenomic analyses (Figure 1).

Under combined condition analysis, *PAPPA* variant rs1895062 (r^2^ to GWAS lead rs1321917 = 0.989) showed a statistically significant allelic effect, with the OA risk alternate “G” allele driving higher reporter activity than the reference allele “A” across donors and barcodes (Fig. 3A). In contrast, 15 other variants at the *PAPPA* locus that were also in high LD (r^2^>0.7) with the GWAS lead exhibited nonsignificant allelic differences (Supplementary Fig. 1). To independently validate the MPRA-identified effect, we cloned the reference and alternate alleles of rs1895062 upstream of the minimal promoter–GFP reporter and quantified reporter activity. Flow cytometric analysis revealed a rightward shift in GFP fluorescence driven by the alternate allele relative to the reference allele in both baseline and stimulated conditions (Fig. 3B). Consistent with this observation, quantification of reporter activity demonstrated a significant increase associated with the oligo harbouring the alternate allele (Fig. 3C). The allelic effect showed a consistent direction across 6 independent donors in both conditions.

**Figure 3.**
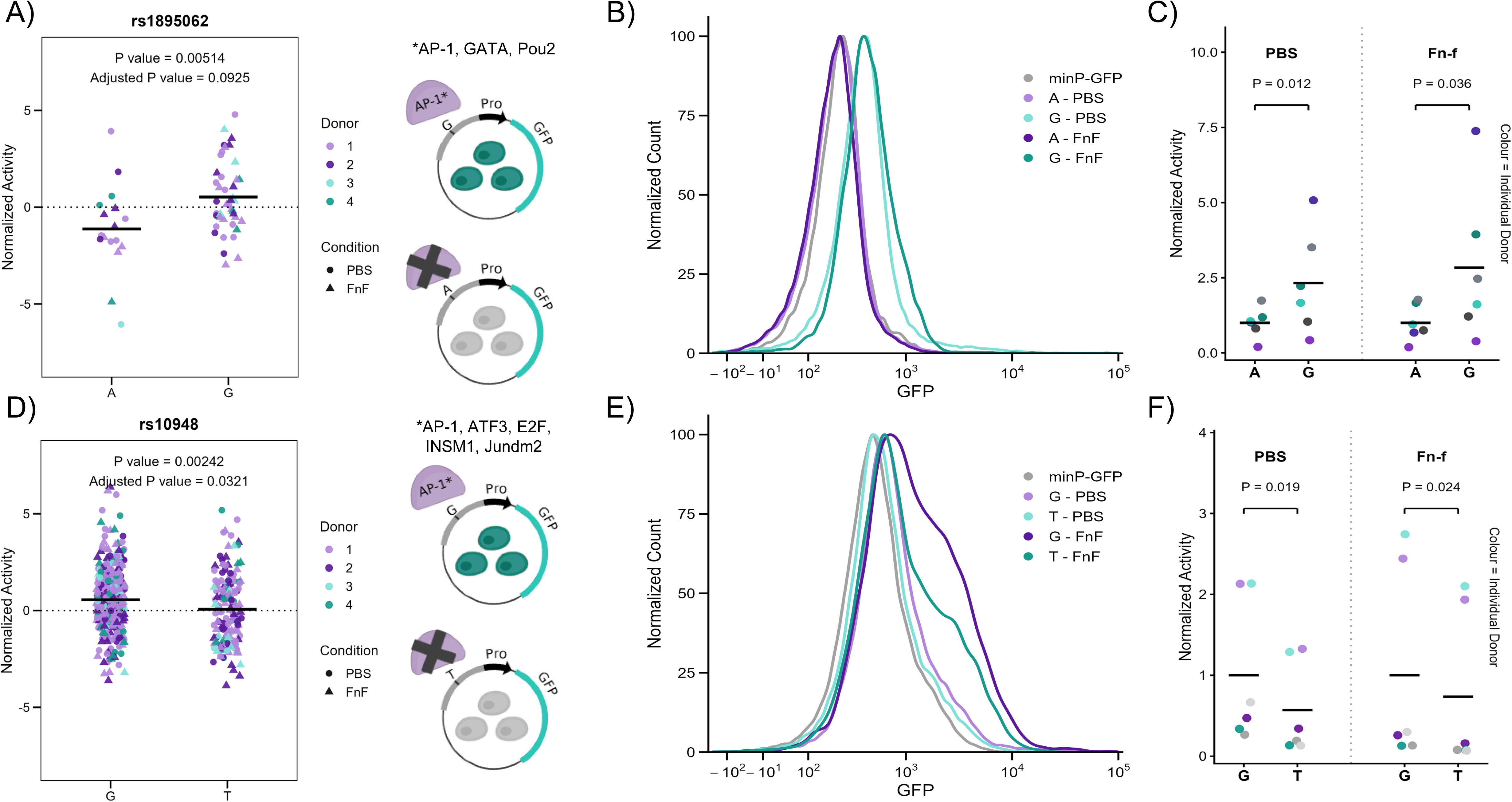
MPRA resolves rs18G5062 as a functional regulatory variant at the *PAPPA* locus and rs10G48 at the SLC44A2 locus. (A) Allele-specific MPRA activity for rs1895062 at the PAPPA locus after combining baseline (PBS) and stimulated (Fn-f) conditions. Each point represents an individual barcode measurement across donors (n = 4). The alternative G allele exhibited significantly increased reporter activity relative to the reference A allele (adjusted p = 0.0925, two-sided t-test following one-pass Grubbs outlier removal). (B) Flow cytometric analysis of GFP reporter constructs carrying the reference or alternate allele of rs1895062 under PBS and Fn-f conditions. Histograms show representative distributions from a single donor. (C) Quantification of the fraction of GFP-high cells for rs1895062 reporter constructs across donors. Points represent individual donor measurements. The alternative G allele significantly increased the fraction of GFP-high cells under both PBS (p = 0.012) and Fn-f (p = 0.036) conditions (paired ratio t-test). (D) Allele-specific MPRA activity for rs10948 at the SLC44A2 locus following combined-condition analysis. Each point represents an individual barcode measurement across donors (n = 4). The reference allele exhibited significantly increased reporter activity relative to the alternate allele (raw p = 0.00242; locus FDR = 0.0321, two-sided t-test following one-pass Grubbs outlier removal). (E) Flow cytometric analysis of GFP reporter constructs containing the reference or alternate allele of rs10948 under PBS and Fn-f conditions. Histograms show representative distributions from a single donor. (F) Quantification of the fraction of GFP-high cells for rs10948 reporter constructs across donors. Points represent individual donor measurements. The reference G allele significantly increased the fraction of GFP-high cells (p = 0.0310, two-sided paired t-test).

We next examined the *SLC44A2* locus, where MPRA identified rs10948 as a candidate regulatory variant in the combined condition. Variant rs10948 exhibited increased reporter activity by MPRA for the OA risk reference “G” allele relative to the alternate “T” allele when analysed across donors and conditions (Fig. 3D). To validate this effect, reporter constructs containing each allele of rs10948 were assessed using the same GFP-based reporter assay. Flow cytometric analysis demonstrated a reproducible shift in GFP fluorescence associated with the reference allele (Fig. 3E), and quantification of the reporter confirmed significantly greater activity with consistent direction of effect across both conditions of 6 independent donors (Fig. 3F).

### Functional annotation of risk loci

While MPRA experimentally identifies variants with allele-specific regulatory activity, orthogonal genomic data can provide additional insight from the native chromatin context to prioritise candidate causal variants at each OA risk locus. To accomplish this, we integrated chromatin accessibility (ATAC-seq), enhancer activity (H3K27ac), and chromatin conformation (Hi-C) in primary OA chondrocytes as well as ChromHMM annotation of publicly available data from vitro differentiated chondrocytes (Supplementary Table S5, Fig. 4A-C).

**Figure 4.**
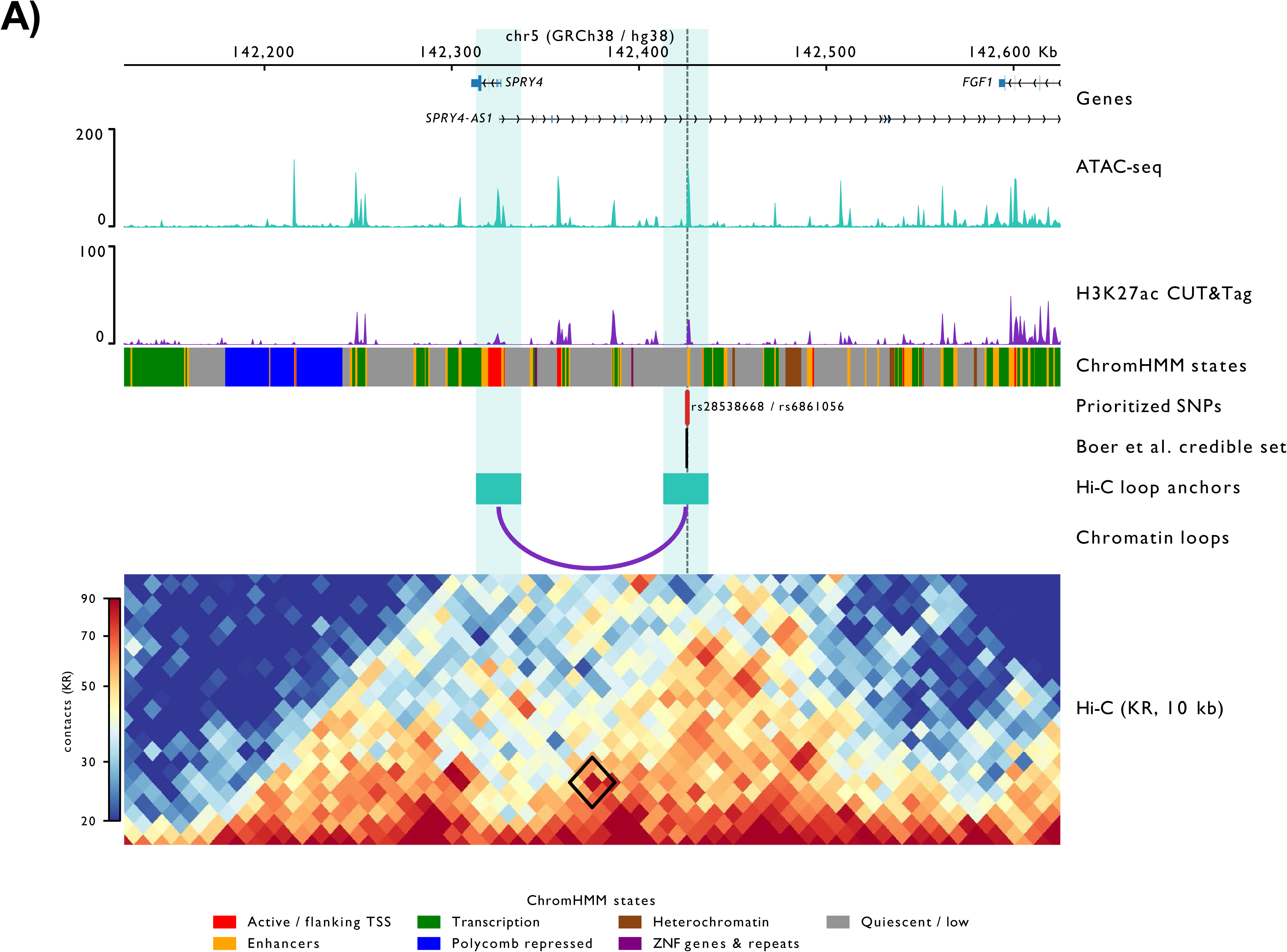
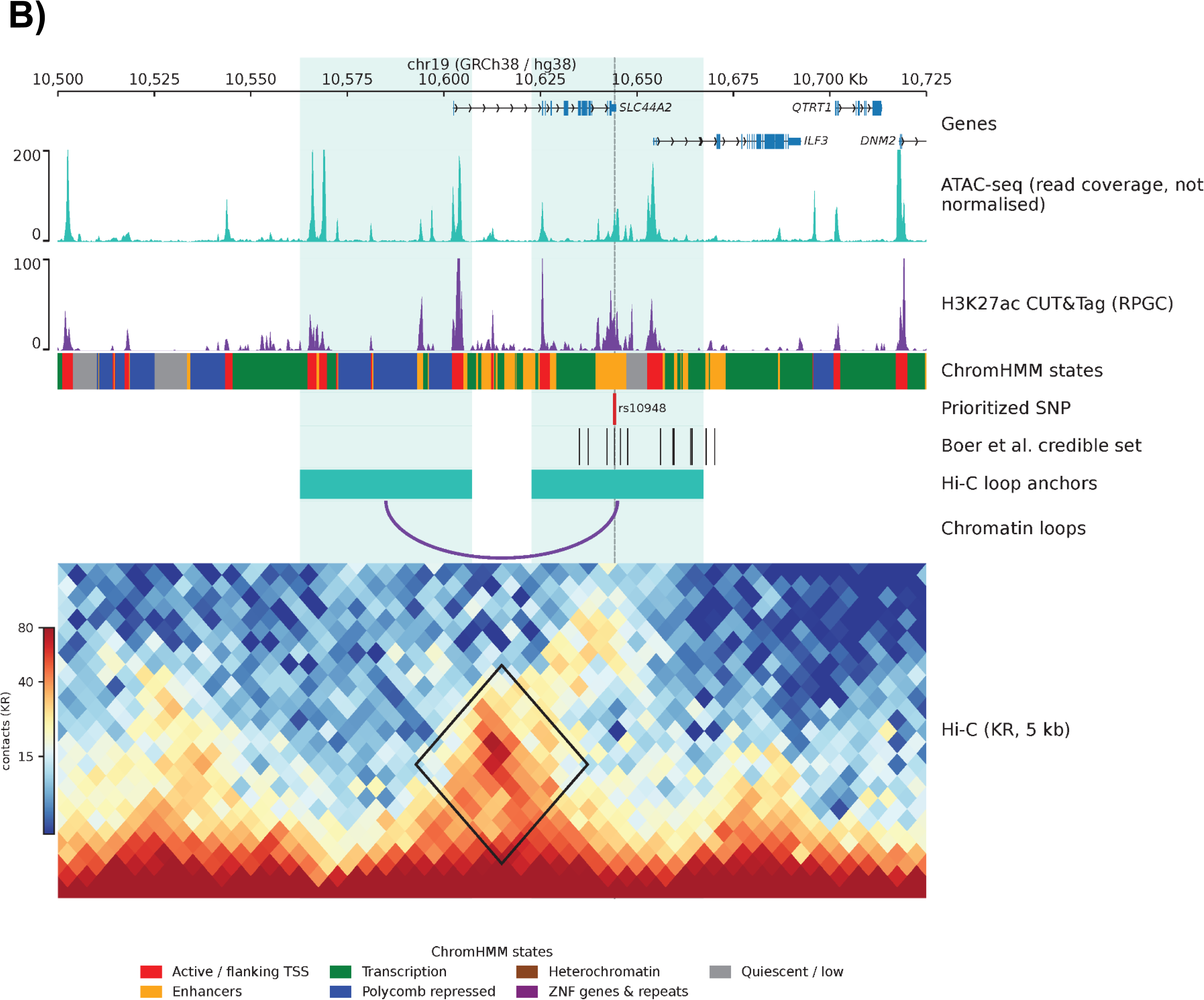
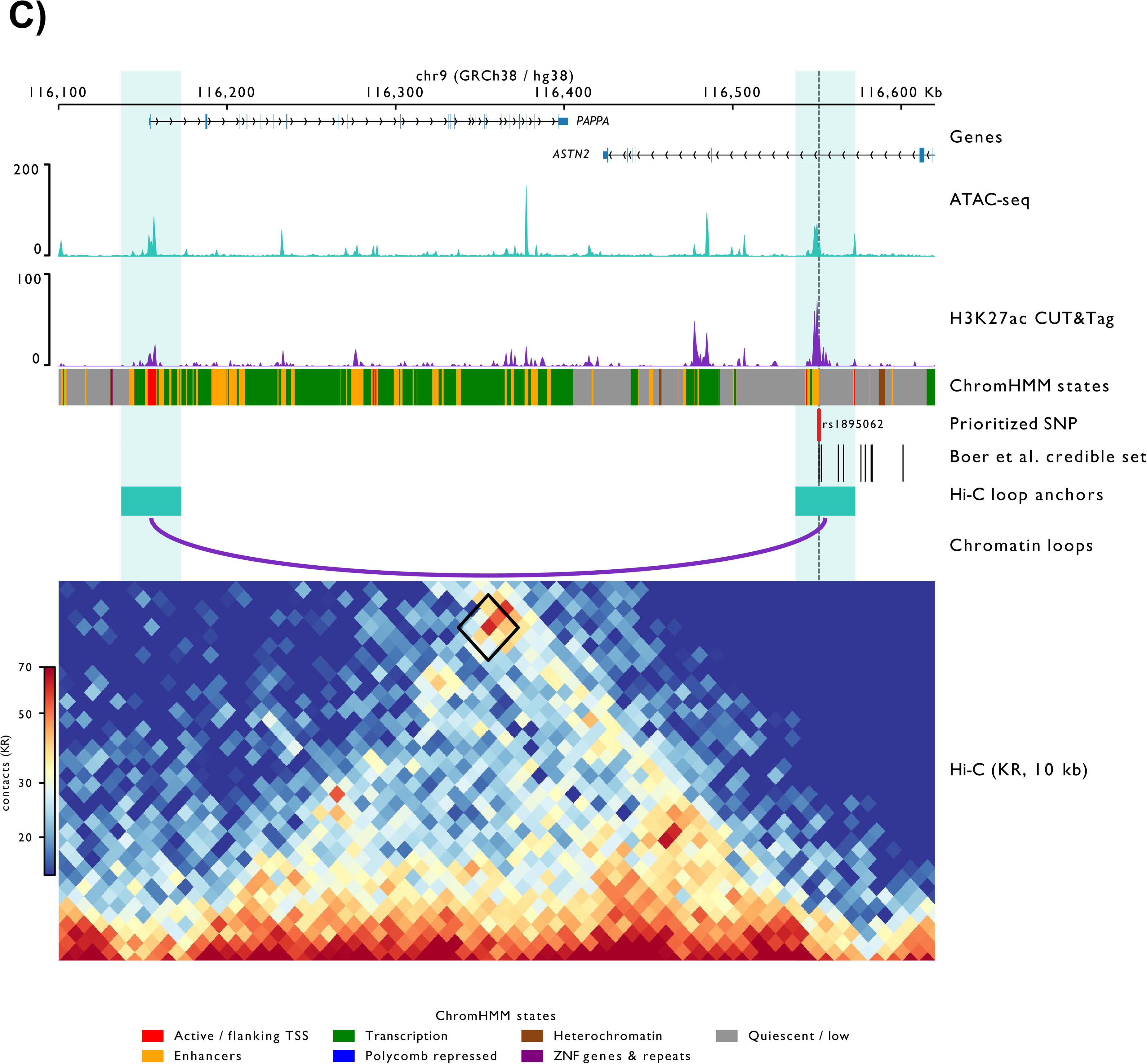
Integration of epigenomic and chromatin interaction data to prioritise candidate regulatory variants at OA risk loci. Genomic landscapes of three OA-associated loci showing the integration of genetic, epigenomic, and chromatin conformation data. Tracks here show gene annotations (top), chromatin accessibility data in primary chondrocytes from OA cartilage (ATAC-seq), active enhancer marks data in primary chondrocytes from OA cartilage (H3K27ac CUT&Tag), ChromHMM-derived chromatin states in *in vitro* differentiated chondrocytes, positions of OA-associated variants in loops with gene promoters (Bittner *et al*.), credible SNP sets at locus of interest from two OA GWAS (Boer *et al*., Hatzikotoulas *et al*.), and Hi-C contact maps from primary chondrocytes (Bittner *et al*.).

Across the three prioritized loci (*PAPPA*, *SLC44A2*, *SPRY4*)^24^, a total of 28 variants were identified within the associated fine-mapped credible sets (Supplementary Table S5). Integration of chromatin accessibility, epigenetic marks consistent with enhancer regions, and chromatin interaction datasets substantially reduced the number of candidate functional variants. Of these 28 variants, only 4 satisfied all three criteria in that they overlap ATAC-seq peaks in primary OA chondrocytes, overlap H3K27ac-marked regulatory regions, and are located within chromatin loop anchors connected to candidate target gene promoters. This progressive refinement highlighted a small subset of variants with multiple independent lines of evidence supporting a regulatory role in disease-relevant cells.

At the *SPRY4* locus (Fig. 4A), multiple variants resided within regions classified as enhancers by ChromHMM and were positioned within chromatin loop anchors. However, only rs28538668 and rs6861056 also demonstrated evidence of both chromatin accessibility and H3K27ac enrichment in primary OA chondrocytes, whereas rs10062749 and rs10076308 lacked support from these additional epigenomic datasets.

Similar refinement was observed at the *SLC44A2* locus (Fig. 4B). Although several variants were annotated as enhancers by ChromHMM, only rs10948 overlapped chromatin accessibility, H3K27ac enrichment, and a chromatin interaction with the candidate target gene. This variant was also identified in our MPRA as an emVar and is a previously reported eQTL^14^, further supporting its potential regulatory role. In contrast, variants such as rs8099926 and rs1975163 were annotated as enhancer elements but lacked evidence of enhancer activity or chromatin looping in primary OA chondrocytes, suggesting that regulatory annotations derived from model systems may overestimate the number of functional variants at a given locus.

At the *PAPPA* locus (Fig. 4C), the lead GWAS variant (rs1321917) showed no evidence of chromatin accessibility, enhancer activity, or chromatin looping in our OA chondrocyte datasets. In contrast, rs1895062 was located within a region displaying all three regulatory features. Notably, rs1895062 was the only variant of the 16 tested in this locus that exhibited allele-dependent regulatory activity in our primary chondrocyte MPRA, resulting in its classification as an emVar. Importantly, this variant was not annotated as regulatory by the ChromHMM based on the only cartilage-related cell type present in the Roadmap Epigenomics Consortium (E049, Mesenchymal Stem Cell Derived Chondrocyte Cultured Cells)^26^; it instead resides within a region classified as quiescent in this cell type. This discrepancy highlights the importance of generating functional genomic maps directly in primary OA tissue, as reliance on public datasets alone would have excluded rs1895062 despite strong regulatory evidence in OA chondrocytes.

### CRISPR-based perturbation of PAPPA and SPRY4 loci confirm regulatory effects

To investigate whether the identified SNPs reside in functional regulatory elements that affect target gene expression, we generated SW1353 chondrosarcoma cell lines stably expressing either dCas9-VPR for CRISPR-based activation (CRISPRa) or dCas9-KRAB for interference (CRISPRi). These cell lines were transduced with lentiviral pools of gRNAs targeting regions within 200 bp of rs1895062 within the *PAPPA* locus, rs10948 within *SLC44A2* locus, and rs6861056 and rs28538668 within the *SPRY4* locus.

Confirming the efficacy of the CRISPRa system, a positive control gRNA targeting the promoter of *IL1RN*, a low abundance gene in the cell line that has often been used as a test locus^27,28^, resulted in a ∼115-fold increase in gene expression (*p* = 0.0210, Supplementary Fig. 2A). Similarly, to validate the CRISPRi-mediated repression system, we targeted the promoter of *RAB1A as a* previously characterised target^28^ and showed a significant reduction in gene expression of 1.8-fold in comparison to non-targeting control (*p* = 0.0011, Supplementary Fig. 2B).

CRISPR-mediated perturbation of the *SPRY4* locus revealed consistent evidence of enhancer activity. Targeting rs6861056 with CRISPRa resulted in a modest but significant upregulation of *SPRY4* expression (*p* = 0.0082, Fig. 5A), whereas CRISPRi produced a stronger effect, with an approximately 2.5-fold reduction in gene expression relative to scrambled control (*p* = 0.0475, Fig. 5A). CRISPRa targeting of a second prioritized SNP in the same locus, rs28538668, did not result in a significant change in *SPRY4* expression. However, CRISPRi perturbation led to a robust decrease in gene expression, comparable to that observed for rs6861056 (P = 0.0071, Fig. 5B). In addition to *SPRY4*, expression of *NR3C1*, the gene proposed by Boer *et al*. as the effector gene at this locus, was also assessed following CRISPR perturbations. No significant changes in *NR3C1* were observed (Supplementary Fig. 3A), further supporting *SPRY4* as the likely effector gene at this locus. Notably, rs6861056 and rs28538668 are separated by only 115 bp, are in very high LD (r^2^=0.97), and reside within the same putative enhancer element, indicating that they likely represent a shared regulatory signal. The observed differences in CRISPRa response are therefore most likely attributable to technical factors, such as gRNA efficiency or positioning relative to key regulatory motifs, rather than reflecting distinct biological functions.

**Figure 5.**
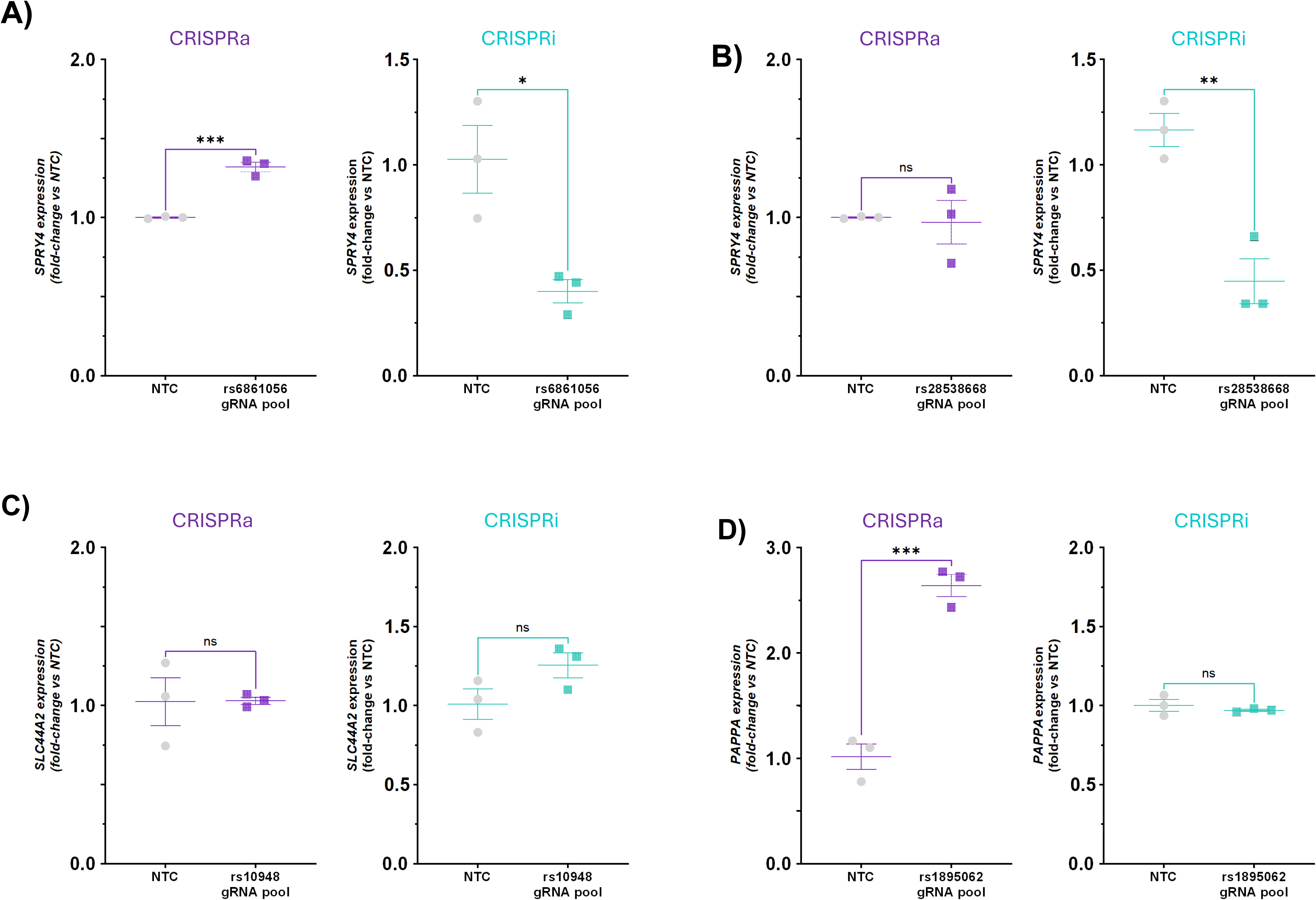
CRISPR-based perturbation of OA-associated loci to functionally test regulatory effects of prioritised loci on predicted target genes. **(A–D)** Quantitative PCR analysis of target gene expression following CRISPR activation (CRISPRa; in purple; left panels) or CRISPR interference (CRISPRi; in teal; right panels) targeting putative enhancer regions containing OA-associated variants. SW1353 cells stably expressing dCas9-VPR (CRISPRa) or dCas9-KRAB-MeCP2 (CRISPRi) were transduced with pooled sgRNAs targeting regions within ±200 bp of each variant. Gene expression is shown relative to non-targeting control sgRNA (SCR), normalised to ACTB using the comparative Ct method. **(A)** SPRY4 locus: targeting rs6861056 results in increased SPRY4 expression following CRISPRa and decreased expression following CRISPRi. **(B)** SPRY4 locus: targeting rs28538668 shows no detectable change with CRISPRa but decreased SPRY4 expression following CRISPRi. **(C)** SLC44A2 locus: targeting rs10948 does not result in a significant change in SLC44A2 expression with either CRISPRa or CRISPRi. **(D)** PAPPA locus: targeting rs1895062 results in increased PAPPA expression following CRISPRa, with no significant change observed following CRISPRi. Bars represent mean ± SEM. Statistical significance was assessed using two-sided Welch’s t-tests, with p-values indicated on each panel.

CRISPR-mediated perturbation of the *SLC44A2* intronic risk locus containing rs10948 did not result in a significant change in gene expression with either CRISPRa or CRISPRi (Fig. 5C). These results do not support a detectable regulatory effect of this element on *SLC44A2* expression under the conditions tested. However, this does not exclude a functional role for rs10948, as regulatory effects may be context-dependent or require specific cellular states, stimuli, or alternative perturbation strategies to be revealed.

Our final perturbation targeting rs1895062 revealed a strong regulatory effect in the distal gene *PAPPA*, located approximately 400 kb away, and previously linked to the SNP via chromatin looping^24,14^. CRISPR-mediated activation of this locus resulted in a robust and significant ∼2-fold increase in *PAPPA* expression (*p* = 0.0006, Fig. 5D), demonstrating that this putative enhancer can drive transcription of the target gene over long genomic distances. In contrast, CRISPR-interference did not result in a significant reduction of *PAPPA* expression (Fig. 5D). This asymmetry between CRISPRa and CRISPRi responses may reflect incomplete repression, redundancy within the local regulatory landscape, or contexts with higher *PAPPA* expression to fully reveal repressive effects.

### Perturbation of the putative PAPPA regulatory element drives transcriptional reprogramming

The CRISPR studies provided evidence for a functional interaction between the enhancer containing rs1895062 and *PAPPA*. Given that this regulatory element was also validated by MPRA and low-throughput reporter assays, we performed RNA-seq on the CRISPRa perturbation experiments to assess the broader transcriptional impact of this regulatory interaction.

Differentially expressed genes (DEGs) were identified by comparing cells subjected to CRISPRa of the PAPPA regulatory region to non-targeting controls. In total, we identified 2,321 significant DEGs (adjusted *p* < 0.05, |log2FC| >1.5), Supplementary Table S6), suggesting substantial and broad transcriptional rewiring (Fig. 6A). Within the topologically associating domain (TAD) containing rs1895062, *PAPPA* was the only gene that met these predefined thresholds of significance and effect size (*p* = 0.0048, log2FC = 1.73), consistent with its identification as the target gene by DNA looping and eQTL analysis. The other two genes within the TAD, *ASTN2* and *TRIM32*, also exhibited statistically significant expression changes, however, neither surpassed the predefined effect-size threshold of |log2FC| >1.5 (Supplementary Fig. 4).

To further characterise the transcriptional response to CRISPRa-mediated enhancer activation, differential expression results were integrated with gene set enrichment analysis (GSEA) using the MSigDB Hallmark gene sets^29^. Several pathways with direct relevance to OA pathogenesis were significantly altered following perturbation of the locus (Fig. 6B). Among upregulated genes, pathways associated with hypoxia and inflammatory signalling, including TNFα signalling via NF-κB, were significantly enriched. In contrast, pathways related to cell-cycle progression, including E2F targets and G2M checkpoint, were significantly depleted. Cholesterol homeostasis was also among the most strongly enriched Hallmark gene sets, indicating broader metabolic changes following enhancer activation.

Consistent with these pathway-level changes, leading-edge genes contributing to the enriched Hallmark gene sets included *HIF1A* and *CAS* within the hypoxia signature, as well as *E2F2* and *PCNA*, key regulators of cell-cycle progression and DNA replication, which were among the most significantly downregulated transcripts (Supplementary Table S6-7). In addition, examination of the most strongly differentially expressed genes identified several biologically relevant changes in extracellular matrix regulation. Matrix remodelling enzymes including *LOX*, *ADAMTS15* and *PLOD2* were upregulated, whereas structural matrix components such as *COL1A1* and *COL5A2* were downregulated. Increased expression of *ILCR* and several components of the NF-κB pathway further supported activation of inflammatory signalling programmes. Enrichment of the cholesterol homeostasis gene set was similarly accompanied by increased expression of multiple genes involved in cholesterol biosynthesis. Together, these findings indicate that activation of this locus is associated with coordinated transcriptional changes affecting inflammatory signalling, hypoxic responses, cellular metabolism, proliferative capacity and extracellular matrix regulation, processes that are highly relevant to cartilage homeostasis and OA pathogenesis.

Finally, we compared the CRISPRa-induced transcriptional response to publicly available transcriptomic data from OA cartilage^30^ to determine whether these transcriptional changes are relevant to disease pathology. We observed substantial overlap in transcriptional changes between the CRISPRa perturbation and gene expression profiles of damaged cartilage compared to lower-grade cartilage from the same OA patients. Several genes exhibited concordant directionality of expression changes across both datasets, including key ECM regulators such as *LOX* and *ADAMTSC*, as well as *PAPPA* itself (Fig. 6C). At the pathway level, GSEA revealed strong concordance between the two datasets, with shared upregulation of pathways associated with hypoxia, upregulation of inflammatory signalling, and cholesterol homeostasis (Fig. 6D). These findings suggest that activation of the *PAPPA* locus recapitulates key transcriptional features of OA cartilage, supporting the functional relevance of this regulatory interaction in disease-associated biological processes.

The results stemming from experimental follow-up of prioritized OA loci are summarized in Figure 7. Three of ten interrogated loci did not produce significant emVars. Of the seven loci that did, two were also prioritized by an independent strategy and were evaluated by both low throughput reporter assays and CRISPR perturbation. The variant rs10948 was an emVar with low throughput validation, but the CRISPR perturbations did not result in a significant change in expression of the *SLC44A2* target gene. Both CRISPR activation and inhibition of the non-coding OA risk region containing rs28538668 and rs6861056 resulted in changes to *SPRY4* gene expression, but this locus was not evaluated by reporter assays. The strongest evidence was for *PAPPA*, as variant rs1895062 was found to be an emVar that validated in low throughput reporter assays and CRISPR activation significantly altered target gene expression.

**Fig.6.**
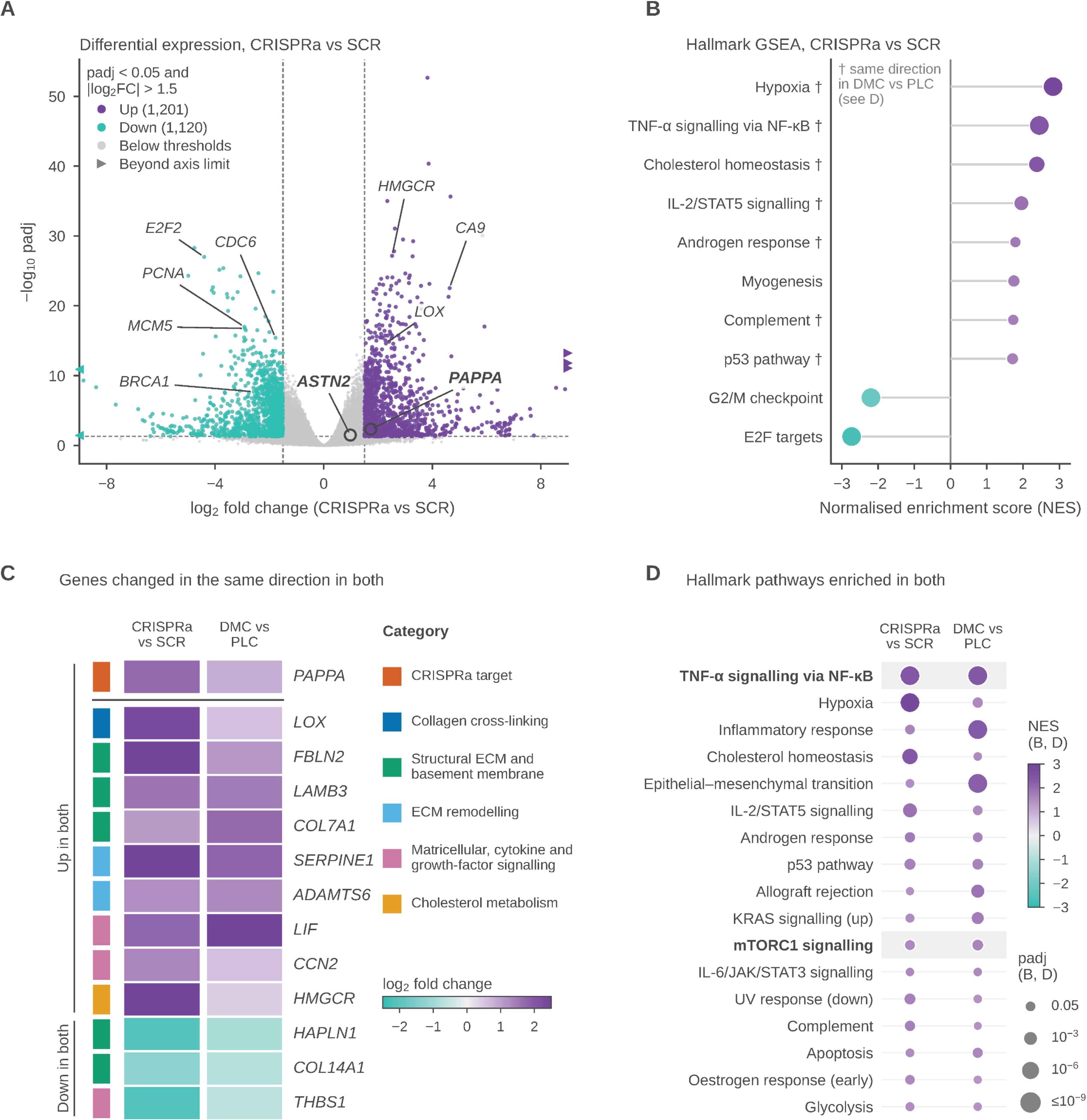
CRISPRa of the rs1895062 enhancer rewires transcription and recapitulates OA cartilage pathway changes. **(A)** Volcano plot of differential expression following CRISPRa of the rs1895062 enhancer versus non-targeting scrambled control (SCR) (Supplementary Table S6). Purple (up, n = 1,201) and teal (down, n = 1,120) points meet padj < 0.05 and |log2FC| > 1.5 (dashed lines); grey points do not. Triangles mark five genes beyond the axis limits (±9). Outlined points are the candidate target genes: *PAPPA* (log2FC = 1.73, padj = 0.005; induction confirmed by RT-qPCR, Fig. 5) and *ASTN2* (log2FC = 0.98, padj = 0.03), significant but below the fold-change threshold. **(B)** Hallmark gene set enrichment analysis (fGSEA), CRISPRa versus SCR. The eight most positively and two most negatively enriched pathways by normalised enrichment score (NES) are shown, of 25 with padj < 0.05 (Supplementary Table S7). Dot colour, NES; dot size, padj (keys beside D). †, enriched in the same direction in damaged medial condyle (DMC) versus preserved lateral condyle (PLC) cartilage (D). **(C)** Genes changed significantly (padj < 0.05) and in the same direction in both comparisons (Supplementary Tables S6, S8). Thirteen of 578 concordant genes are shown, selected for published links to IGF signalling, extracellular matrix and chondrocyte biology. *PAPPA*, the CRISPRa target, is shown above the line; remaining genes are grouped by direction and category, then ordered by mean log2FC. Colour, log2FC (capped at ±2.5). **(D)** All 17 Hallmark pathways with padj < 0.05 and concordant direction in both comparisons, ordered by mean NES (Supplementary Table S9). Shading marks pathways discussed in the text: TNF-α signalling via NF-κB, and mTORC1 signalling, downstream of the *PAPPA*-regulated IGF axis.

**Figure 7.**
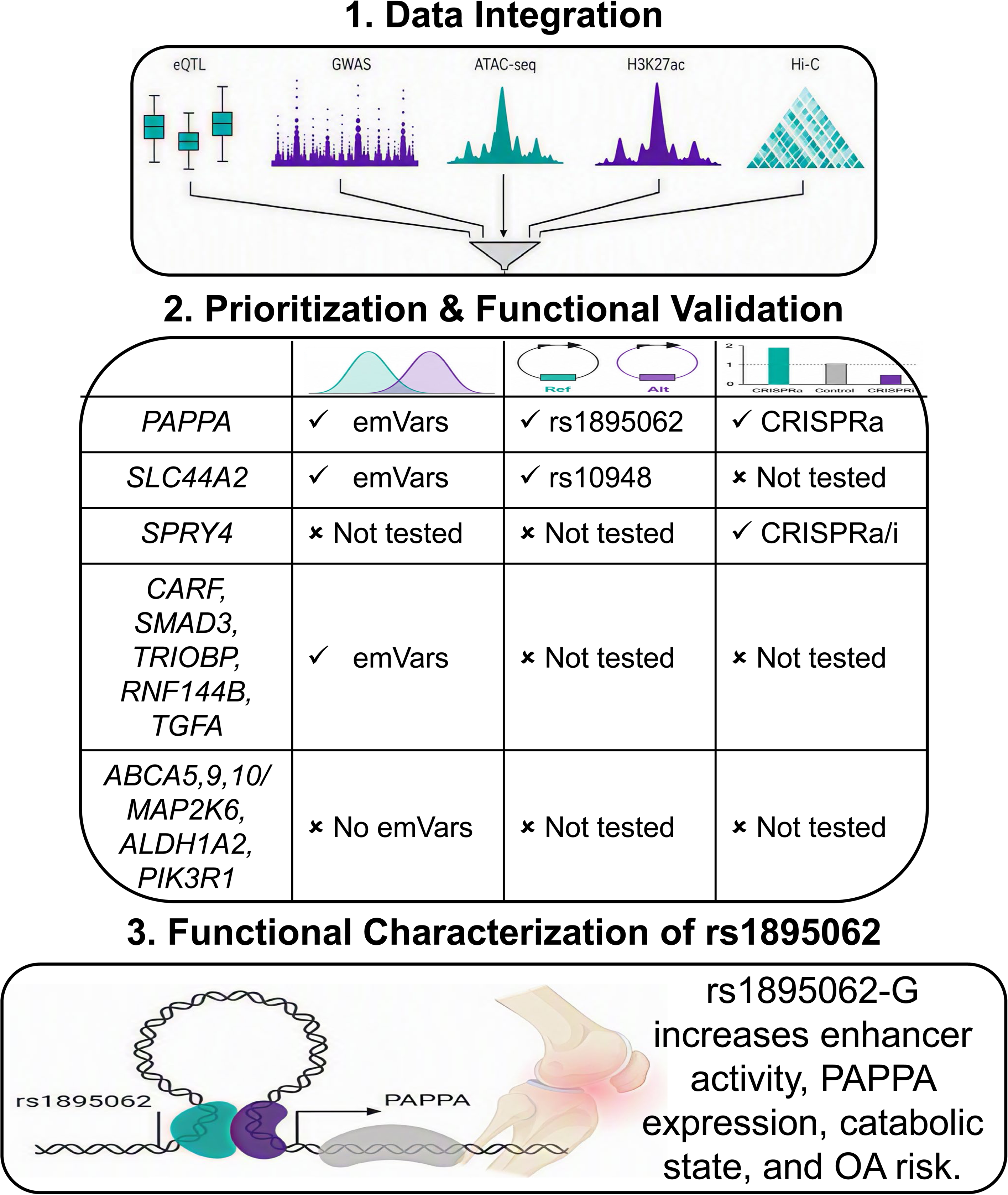
Integrative functional characterization of OA-associated genetic loci. Complementary genetic, epigenomic, and functional approaches were integrated to characterize regulatory mechanisms at OA risk loci. MPRA identified variants with allele-specific regulatory activity, with selected emVars validated by low-throughput reporter assays, while chromatin accessibility, H3K27ac enrichment, and chromatin conformation provided complementary information about the regulatory landscape and potential target genes. At the *PAPPA* locus, integration of these data supports a model linking OA-associated regulatory variation to *PAPPA*, illustrating how complementary approaches can move from genetic association toward functional interpretation of OA risk loci.

## Discussion

Despite large advances in our understanding of OA genetics, there are currently no approved disease-modifying therapies for the disorder. This highlights an unmet clinical need for approaches that translate genetic associations into target genes and molecular mechanisms, thereby improving our understanding of disease pathophysiology and facilitating the identification of relevant therapeutic strategies^31^. Combining multiple complementary lines of evidence is the most robust framework to resolve GWAS signals, allowing for causal variants, effector genes, and downstream molecular pathways to be identified with greater confidence. Through this approach, we sought to systematically test the enhancer activity of candidate regulatory sequences harbouring OA-associated variants, utilize disease-relevant multi-omic integration, and employ CRISPR-based perturbation to resolve OA GWAS signals and improve the mechanistic interpretation of complex disease genetics.

We demonstrate that MPRAs performed in primary human chondrocytes can robustly identify allele-dependent regulatory activity at OA-associated loci. By first establishing assay performance and concordance with orthogonal reporter measurements, and then applying this framework to genetically complex regions, we show that MPRA can resolve functional regulatory variants within disease-relevant primary cells. Importantly, while multiple GWAS-linked variants were evaluated across OA-associated loci, only a subset demonstrated detectable regulatory effects in this experimental context, highlighting the value of functional screening approaches for refining candidate causal variants within linkage disequilibrium blocks. While other promising candidates were identified and could be evaluated in future experiments, we prioritized performing low-throughput reporter assays for rs1895062 at the *PAPPA* locus and rs10948 at the *SLC44A2* locus due to the complementary evidence provided by integrated genomics analysis. Both MPRA and low-throughput assays confirmed that the OA risk allele at rs1895062 (G, associated with higher *PAPPA* expression by eQTL) results in higher enhancer activity. Similarly, we confirmed that the OA risk allele at rs10948 (G, associated with higher *SLC44A2* expression by eQTL) results in higher enhancer activity. Together, these results indicate that MPRAs performed in primary human chondrocytes can distinguish functional regulatory variants within OA-associated loci and identify reproducible allele-dependent enhancer activity for prioritized candidate variants.

The integration of primary OA chondrocyte epigenomic data with chromatin interaction maps enabled substantial refinement of candidate causal variants and effector genes at each locus. While previous analyses identified enhancer–promoter interactions and generated plausible variant-to-gene hypotheses^31^, incorporation of disease-relevant chromatin accessibility and enhancer activity data prioritized a smaller subset of variants with direct evidence of regulatory potential. Our data highlight how disease-relevant regulatory landscapes can differ from broadly available datasets. For example, variant rs1895062 was classified as quiescent by ChromHMM of public data but was found to be within an accessible and active chromatin region in our OA chondrocyte datasets. Furthermore, chromatin organisation is particularly helpful for assigning effector genes at large distances with high confidence. This is critical, as our previous chondrocyte eQTL study demonstrated that more than half of signals were matched to eGenes further than at least one other gene^14^. These analyses highlighted rs1895062 (*PAPPA*), rs10948 (*SLC44A2*), rs6861056 (*SPRY4*), and rs28538668 (*SPRY4*) as the strongest candidates for functional interrogation and formed the basis for subsequent CRISPR-mediated enhancer perturbation experiments. Importantly, the prioritisation strategy described here was based on the largest OA GWAS available at the time, performed by Boer *et al.*^7^. Since then, a substantially larger meta-analysis has identified many additional OA-associated signals and further refined several previously reported loci. Of note, all four variants prioritised in the present study (rs1895062, rs10948, rs6861056 and rs28538668) remain within their respective credible sets in the updated analysis. At the *PAPPA* locus, fine-mapping in this highly powered GWAS has reduced the credible set to only two candidate variants, one of which is rs1895062, providing additional support for its prioritisation as the likely functional variant underlying this association.

We show the power in using bidirectional CRISPR-based perturbations to functionally interrogate non-coding regulatory elements. The complementary use of CRISPR-activation and interference allowed us to assess enhancer activity across loci with different basal levels of gene expression, providing a dynamic framework to capture both activating and repressive regulatory effects. Across the loci interrogated in this study, we observed a spectrum of regulatory effects following CRISPR-based perturbation. The regulatory element containing rs1895062 showed an effect on *PAPPA* expression in the CRISPRa context but not with CRISPRi. In contrast, *SPRY4* showed the strongest effects in response to CRISPRi, although rs6861056 also showed an effect with CRISPRa. *SLC44A2* expression was not significantly affected by either CRISPRa or CRISPRi of the prioritized regulatory element. These examples demonstrate that the CRISPRa/CRISPRi approach provides an effective strategy to characterise functional consequences of non-coding risk SNPs.

The *SLC44A2* locus provided an example where multiple lines of evidence supported regulatory potential, yet CRISPR perturbation failed to produce measurable changes in the predicted target gene. While rs10948 demonstrated reproducible allele-dependent enhancer activity in both MPRA and low-throughput reporter assays, neither CRISPRa nor CRISPRi significantly altered *SLC44A2* expression. Several explanations may account for this observation. Regulatory activity at this locus may be highly context dependent, requiring environmental or inflammatory stimuli not present under the experimental conditions used here. Consistent with this possibility, rs10948 was previously identified as an eQTL in Fn-f treated chondrocytes but not under basal conditions^14^. Although the precise mechanism remains unresolved, these findings highlight the challenges associated with translating chromatin interaction maps and enhancer activity into experimentally demonstrable gene regulatory effects. Nevertheless, *SLC44A2* remains a biologically plausible OA candidate. The gene encodes a choline transporter involved in phospholipid metabolism and mitochondrial function^32^, processes that have been implicated in OA pathogenesis^33^, and *SLC44A2* expression is altered with OA in both cartilage and subchondral bone^34^.

Perturbation of the *SPRY4* locus revealed a complex but ultimately informative regulatory architecture. Integration of chromatin accessibility, enhancer activity and chromatin interaction data prioritised rs6861056 and rs28538668 within a putative enhancer physically interacting with the *SPRY4* promoter. Functional perturbation experiments consistently supported *SPRY4* as the principal effector gene at this locus. Repression of either regulatory element by CRISPRi resulted in reduced *SPRY4* expression, whereas expression of the neighbouring gene *NR3C1* remained unchanged, providing experimental support for the enhancer-promoter interactions predicted by our multi-omic analyses. In contrast, transcriptional activation produced more modest effects, with only one of the two targeted regions generating a detectable increase in SPRY4 expression. The biological functions of *SPRY4* further support its candidacy as an OA effector gene. *SPRY4* is a negative regulator of receptor tyrosine kinase and MAPK/ERK signalling, and expression of this gene has been associated with protection from OA-related phenotypes including chondrocyte hypertrophy, senescence and extracellular matrix degradation^35^. Together with the functional perturbation data presented here, these findings support altered *SPRY4* regulation as a plausible mechanism linking genetic variation at this locus to OA susceptibility.

*PAPPA* is an interesting locus in the context of OA, as previous investigation showed that *PAPPA* expression increases with age, OA, and in response to the Fn-f stimulus^14^. *PAPPA* encodes a secreted metalloproteinase that regulates the bioavailability of insulin-growth factor (IGF) through cleavage of specific IGF binding proteins. Given the well-established role of IGF signalling in cartilage homeostasis and repair, dysregulation of this pathway has been implicated in the pathogenesis of OA^36,37^. Higher levels of *PAPPA* are implicated in senescence, as inducing senescence through loss of sirtuins alters DNA looping to the *PAPPA* promoter and leads to higher expression^38^. Consistent with a critical role for *PAPPA*, our work shows that CRISPRa of the regulatory region containing rs1895062 coordinated changes across multiple biological pathways relevant to cartilage biology. We observed upregulation of numerous genes associated with ECM remodelling, inflammatory, and hypoxia-associated signalling programmes, whereas there was an overall suppression of cell-cycle and proliferation related processes. These observations are in line with known pathogenic pathways in OA, where dysregulated ECM turnover, chronic inflammation, and altered chondrocyte proliferation contribute to disease progression^39,40^. Importantly, comparison of our CRISPRa perturbed cells with publicly available transcriptomic data from OA cartilage showed significant concordance at both gene and pathway levels, suggesting that activation of the rs1895062 enhancer recapitulates key features of the OA transcriptional landscape. In addition to these well-described OA pathways, we also identified strong enrichment of cholesterol homeostasis pathways following CRISPRa perturbation, which may be related to a role for PAPPA-mediated signalling in broader cellular metabolic states as has been shown in other tissue types^41^.

Collectively, these studies provide a disease-relevant framework for resolving enhancer-promoter interactions and identifying the causal regulatory variants and target genes that underlie OA susceptibility. However, there are important limitations that should be considered. First, our prioritization based on eQTL, chromatin conformation, and genomics integration leave many other OA GWAS loci unexplained. The insight gained from the current investigations will enable larger scale MPRAs, genomic integration, and CRISPR perturbations to identify more specific examples and broader themes of how genetic risk operates in OA. Second, the CRISPR experiments were carried out in a chondrosarcoma-derived cell line, which, although widely used^42,43^, may not fully recapitulate the transcriptional and regulatory landscape of primary chondrocytes *in vivo*. While integration of multi-omics and publicly available OA transcriptomics data from primary OA patients provides support for disease relevance, further functional validation in primary cells or *in vivo* systems will be required to fully establish the role of these regulatory interactions in OA pathogenesis. Third, additional work will be required to fully define the molecular mechanisms linking the prioritised regulatory variants to OA pathogenesis. While *PAPPA* emerged as the strongest candidate effector gene in the current study, the incomplete CRISPR validation observed at the *SLC44A2* locus highlights the context-dependent nature of regulatory interactions and the need for future studies in alternative cellular and disease-relevant stimulation conditions. At a mechanistic level, it will be important to determine whether allele-specific differences influence transcription factor binding, as Bittner *et al.* predicted computationally that rs1895062 may affect *AP-1*, *GATA1* and *Pou2*. At a functional level, modulating *PAPPA* in contexts where cell-matrix interactions are important, such as tissue explants or animal models, will be important to further define the role of *PAPPA* in OA pathogenesis and provide a foundation for future therapeutic targeting strategies.

## Conclusions

This work provides direct functional evidence supporting regulatory mechanisms underlying OA genetic risk and establishes a scalable framework for prioritizing noncoding variants for downstream mechanistic investigation in disease-relevant cellular systems. Multiple layers of evidence propose that rs1895062 genotype controls the strength of a regulatory element that alters *PAPPA* gene expression, with activation of this regulatory element leading to widespread transcriptomic change.

## Methods

### MPRA Library

To enable functional assessment of candidate variants, we developed an MPRA library design strategy to generate 200 bp oligos. The final MPRA library consisted of n=832 test variants and n=300 controls. Tested variant selection was based on colocalized primary human chondrocyte eQTLs and OA GWAS signals reported in our previous work which includes 13 colocalized eGenes at 10 genomic loci^14^. For each of these 10 loci, we selected lead variants and all variants in high LD using TopLD^44^ (Population: European, LD threshold: 0.7, and MAF threshold: 1%) resulting in 10 lead SNPs, 355 high LD-linked SNPs, 23 LD-linked indels, and 5 additional SNPs of interest. For each SNP, we designed two oligos, containing either the risk or non-risk allele centred within the oligo sequence using the hg38 reference (n=740). For each indel, we designed four oligos, using a positional shifting strategy to address any bias introduced by the additional flanking sequence in the deleted allele (n=92).

We included three types of controls: (1) 100 ‘active control’ sequences that are likely to induce transcriptional activation, (2) 100 ‘inactive scrambled control’ sequences that are scrambled versions of the active sequences and therefore unlikely to induce transcriptional activation, and (3) 100 ‘emVar control’ sequences which are 50 pairs of identical sequences that differ by a single allele and are expected to exhibit differential regulatory activity. To select sequences for these three control groups, we leveraged publicly available MPRA data on HEPG2, SK-N-H, and K562 cells from Supplementary Table 4 of Siraj et al. 2026,^25^ as well as our previously published chromatin accessibility data in primary human chondrocytes^14^. Additional details on control sequence selection are provided in Supplementary Methods and the full list of oligo sequences is in Supplementary Table 1.

MPRA libraries were constructed using the Tewhey Lab MPRA v3 framework^16^. Barcode–oligo associations were established by high-throughput sequencing prior to cellular assays.

### Normal chondrocyte isolation/culture and MPRA transfections

Primary human articular chondrocytes were isolated from normal cartilage tissue obtained from cadaveric tissue donors as previously described^14^. Use of de-identified donor cartilage was determined not to constitute human subjects research by the University of North Carolina Institutional Review Board. Briefly, chondrocytes were isolated by sequential pronase (Sigma Aldrich) and collagenase (Sigma Aldrich) digestion and cultured in DMEM/F-12 (Thermo Fisher Scientific) supplemented with 10% fetal bovine serum (FBS; Thermo Fisher Scientific). In preparation for nucleofection, cells were switched to antibiotic-free medium. Cells were detached using 0.05% trypsin-EDTA (Thermo Fisher Scientific), washed with PBS (Thermo Fisher Scientific), and pelleted at 300 × g for 5 minutes. Five million cells were resuspended in 100 µL of supplemented P3 Primary Cell 4D-Nucleofector™ X Solution (Lonza Walkersville P3 Primary Cell 4D-Nucleofector X Kit L; V4XP3024). For each reaction, 5 µg of mpra:gfp plasmid library DNA was added and cells were nucleofected using the 4D-Nucleofector™ X Unit (program ER-100).

Following nucleofection, cells were seeded at a density of 2.6 × 10⁵ cells/cm². Medium was replaced after 24 hours with fresh antibiotic-free medium. At 52 hours post-nucleofection, cells were serum-starved, and at 54 hours cells were treated with PBS or 1 µM purified human recombinant fibronectin fragment (Fn-f) in serum-free, antibiotic-free medium. Fn-f is a bioactive matrikine that provides an OA-relevant inflammatory stimulus to elicit transcriptional responses that may not be present in baseline conditions^14,22^. Cells were harvested at 72 hours post-nucleofection using 0.05% trypsin-EDTA for downstream RNA isolation.

Cells were harvested for RNA isolation, GFP mRNA pulldown, cDNA synthesis, and reporter transcript abundance was quantified by barcode sequencing. Details are provided in the Supplementary methods.

### MPRA analysis

Barcode-level RNA and DNA counts were processed using the MPRAsnakeflow pipeline, followed by downstream analysis using a custom R workflow. One donor had low coverage of barcodes across most variants and was excluded, leaving four donors for analysis. Barcode activity was calculated as the log₂ ratio of RNA to DNA counts and median-centred within each donor × condition using scrambled negative-control oligos. Allelic effects were assessed by pairing reference and alternate sequences for each variant and comparing the barcode-level activity distributions between alleles. Multiple testing correction was performed using the Benjamini–Hochberg procedure. Details on the analyses are provided in the Supplementary Methods and the analysis pipeline is available through GitHub (https://github.com/srstuppy/MPRA).

### Low throughput reporter assays

Individual 200 bp oligonucleotides warranting additional investigation were synthesized as double-stranded DNA gene fragments (Twist Bioscience) containing Gibson homology arms and inserted into pMPRAv3:minP-GFP, which was a gift from Pardis Sabeti C Ryan Tewhey (Addgene plasmid # 109036; http://n2t.net/addgene:109036; RRID:Addgene_109036)^16^. Sequence-verified plasmids were expanded and purified using Zymo maxiprep kits for downstream nucleofection. Low-throughput reporter assays were performed in primary human chondrocytes using the same isolation, nucleofection, and treatment conditions described for MPRA transfections. Reporter activity was assessed by flow cytometry and immuno fluorescence (IF) using both native and antibody-based detection of GFP. For flow cytometry, stained cells were analysed to quantify the percentage of GFP-positive cells and/or GFP fluorescence intensity using standard gating strategies, with DAPI used to gate nucleated events. For IF, stained cells were imaged under identical acquisition settings across conditions, and GFP fluorescence was quantified per cell using CellProfiler.

### OA chondrocyte isolation for genomics studies

Human articular cartilage samples were obtained from patients undergoing total knee arthroplasty (TKA) surgery for end-stage OA at the Manchester Orthopaedic Centre, Trafford General Hospital. Ethical approval for the collection and use of human tissue was granted by the North West – Greater Manchester West Research Ethics Committee (REC reference: 23/NW/0136; IRAS project ID: 328269). All participants provided written informed consent prior to tissue collection.

Patient eligibility was determined solely by the ability to provide informed consent. Consequently, individuals unable to independently provide informed consent were excluded from recruitment. No additional exclusion criteria relating to patient demographics or clinical characteristics were applied.

Articular cartilage was dissected from all available anatomical regions of the knee joint and macroscopically graded as either high-grade or low-grade cartilage as defined by the International Cartilage Repair Society^45^. Low-grade cartilage (grade 0-1) were cut into small pieces, washed twice with phosphate-buffered saline (PBS), and set to digest in Dulbecco’s Modified Eagle Medium (DMEM) with 10 mg/mL collagenase II (Thermo Fisher Scientific, 17101015) on a 37°C shaker for 4 hours. Tissue suspensions were then passed through a 70 μm cell strainer and a suspension of single chondrocytes was collected. Cells were washed with PBS, frozen in Recovery Cell Culture Freezing Media (Thermo Fisher Scientific, 12648010) at approximately 1 million cells per aliquot and stored at -150°C for future experiments.

### ATAC-seq

ATAC-seq libraries (n=3 patients) were prepared according to the Omni-ATAC protocol^46^, using 50,000 cells per library and with the following modifications: cell lysis time was increased to 5 minutes, and the duration of the transposition step was extended to 40 minutes. Library size and nucleosome integrity was examined with the Tapestation 4200. Amplified libraries were sequenced on the NovaSeq X Plus platform (Illumina) at a target depth of 50 million reads per library (150bp paired-end reads).

ATAC-seq reads were quality filtered, trimmed, and adapters were removed using fastp v0.23.4^47^. Trimmed reads were mapped to reference genome using bowtie2 v2.5.1^48^, using the very-sensitive pre-set. Duplicate reads were removed using the MarkDuplicates functionality of Picard tools (https://broadinstitute.github.io/picard/). Mitochondrial reads and low-quality alignments (MAPQ<30) were also removed, keeping only properly aligned read pairs. The bamCoverage functionality from DeepTools2 was used to generate coverages from cleaned alignments and the macs3 v 3.0.3 programme was used for peak calling^49^. The ATAC-seq processing pipeline is available on GitHub (https://github.com/jakebutler98/UoM_ATAC_processor).

### CUTGTag

CUTCTag library (n=1 patient) was prepared using the CUTANA CUTCTag Kit (EpiCypher, 14-1102-24s3), according to manufacturer’s protocol. Briefly, 100,000 cells per library were immobilized using kit-provided ConA beads and nuclei were extracted, incubated with 0.5 µg of primary antibody against H3K27ac (EpiCypher, 13-0059) overnight at 4°C. Secondary antibody was added and DNA was tagmented as per kit’s protocol. Resulting libraries were cleaned using AMPure XP beads and library quality was examined with the Tapestation 4200. Amplified libraries were sequenced on the NovaSeq X Plus platform (Illumina) at a target depth of 8 million reads per library (150 bp paired-end reads).

CUTCTag reads were quality-filtered, trimmed, and adapter sequences were removed using fastp v0.23.4^47^. Trimmed reads were aligned to the reference genome using Bowtie2 v2.5.5^48^ in end-to-end mode with the --very-sensitive, --no-mixed, --no-discordant, --phred33, and - -dovetail options. SAMtools v1.23.1^50^ was used to remove mitochondrial reads, alternate contigs, unmapped reads, and low-quality alignments (MAPQ < 2), retaining only properly paired alignments. Additionally, ENCODE blacklist regions were removed using BEDTools v2.31.1^51^. Coverage tracks were generated from the filtered alignments using the bamCoverage function from DeepTools2^52^, and peak calling was performed with MACS3 v3.0.4^49^. The CUTCTag processing pipeline is available on GitHub (https://github.com/CarlosRangel23/CutTagFlow).

### Functional Annotation of OA GWAS Variants

Candidate OA risk loci from Bittner *et al.* were functionally annotated through integration of genetic, epigenomic, and chromatin conformation datasets (Supplementary Table 5).

Briefly, credible SNP sets (posterior probability ≥ 0.03) for each locus were obtained from Boer *et al.*^7^, formatted as BED files, and initially visualised using the Integrative Genomics Viewer (IGV)^53^. Publicly available epigenomic datasets from in vitro differentiated chondrocytes were downloaded from the ENCODE Consortium^54^, including H3K27ac (ENCSR925IAP), H3K4me1 (ENCSR117SLI), and DNase-seq (ENCSR970DQR), alongside ChromHMM-derived regulatory state annotations from the Roadmap Epigenomics Consortium^26^.

To generate disease-relevant regulatory maps, in-house datasets generated from primary OA chondrocytes, including ATAC-seq and CUTCTag profiling of H3K27ac, were analysed in parallel with public datasets. Chromatin interaction data were obtained from Bittner *et al*.^24^, including Hi-C contact matrices (.hic files) and associated loop and anchor annotations derived from supplementary datasets.

For final visualisation, all datasets (genetic variants, epigenomic tracks, and chromatin interaction data) were integrated and plotted using pyGenomeTracks. Hi-C matrices were rendered alongside loop and anchor annotations, and overlaid with epigenomic signal tracks and gene annotations, enabling comprehensive contextualisation of candidate enhancer–promoter interactions at each locus.

### CRISPRa/CRISPRi gRNA Design and Cloning

Guide RNAs (gRNAs) targeting prioritized putative enhancer regions containing SNPs of interest were designed using the CRISPOR tool^55^, focusing on DNA regions 200 bp upstream and 200 bp downstream of OA risk SNPs. For CRISPR activation and interference experiments, we used a pool of three gRNAs targeting regions upstream and downstream of the SNP of interest to maximise targeting efficiency. Forward and reverse oligonucleotide sequences used for cloning CRISPR guide RNAs targeting putative OA-associated enhancer regions at four candidate loci: rs28538668 (*SPRY4* locus), rs6861056 (*SPRY4* locus), rs1895062 (*PAPPA* locus), and rs10948 (*SLC44A2* locus) are listed in Supplementary Table 10.

Complementary gRNA oligo pairs with 5′ CACC (forward) or 5’ CAAA (reverse) overhangs (Thermo Fisher Scientific, 100 mM) were phosphorylated and annealed in a thermocycler (37°C for 30 min; 95°C for 5 min; ramp down to 25°C at 5°C/min) using T4 Polynucleotide Kinase (NEB, M0201S).

For cloning, an in-house generated lentiviral vector plasmid, pLKO5.sgRNA.NeoR, was used. This plasmid was previously adapted from pLKO5.sgRNA.EFS.GFP (Addgene, #57822) by replacing the GFP gene with a neomycin resistance gene. The plasmid was digested with the FastDigest restriction enzyme Esp3I (Thermo Fisher Scientific, ER0451) and FastAP Thermosensitive Alkaline Phosphatase (Thermo Fisher Scientific, EF0654) at 37°C for 2 hours, followed by ligation with the annealed gRNA duplex using T4 DNA Ligase (NEB, M0202S) at 16°C overnight.

The resulting gRNA-containing lentiGuide plasmids were transformed into One Shot™ Stbl3™ Chemically Competent E. coli (Thermo Fisher, C7373-03) by heat shock (42°C for 45 seconds) according to the manufacturer’s instructions. Plasmid DNA was then isolated using the QIAprep Spin Miniprep Kit (QIAGEN, 27104) and Sanger sequenced to confirm insertion of the desired gRNA sequences.

To produce gRNA-containing lentiviruses, a third-generation system was used, requiring four separate plasmids: two packaging plasmids (pRRE, Addgene #12251; pRSV-REV, Addgene #12253), one envelope plasmid (pMD2.G, Addgene #2259), and one transfer plasmid containing the pool of gRNA plasmids targeting the SNP of interest. These plasmids were combined at a mass ratio of 1:2:1:4 and transfected into HEK293T cells. Lentiviral particles were harvested from the supernatant 48 hours post-transfection and stored at −80°C.

### Perturbation of Putative Enhancer Regions with dCASG-VPR and dCASG-KRAB

The chondrosarcoma cell line SW1353 was transduced with lentiviruses encoding either dCas9-VPR-Puro (Addgene, #99373) or dCas9-KRAB-MeCP2 (Addgene, #110821) to generate stable CRISPR activation (SW1353-CRISPRa) and CRISPR interference (SW1353-CRISPRi) cell lines, respectively.

Successfully transduced cells were selected using puromycin and subsequently transduced with gRNA-containing lentiviral particles targeting the genomic regions of interest. Transductions of gRNA lentiviruses were done in triplicates and positive and negative controls were ran simultaneously. Positive controls were targeted to *IL1RN* and *RAB1A* for CRISPRa and CRISPRi, respectively. Negative controls for both perturbations were carried out with a scrambled, non-targeting sequence.

Stable gRNA-expressing SW1353-CRISPRa and SW1353-CRISPRi cells were then selected using neomycin and allowed to recover for one week. Following recovery, cells were lysed and RNA was extracted using the RNeasy Mini Kit (QIAGEN, 74104) for downstream analysis.

### qPCR

Total RNA isolated from SW1353-CRISPRa and SW1353-CRISPRi was DNaseI-digested and reverse transcribed using the SuperScript IV VILO Master Mix (Thermo Fisher Scientific, 11766500) following manufacturer’s instructions. Quantitative PCR (qPCR) was performed using the Luna Universal qPCR Master Mix (NEB, M3003L) with previously validated primers for gene targets of interest^21–23^ and using cycling conditions recommended by the manufacturer. Relative gene expression was calculated using the comparative Ct (2^−ΔΔCt) method^24^, with *ACTB* used as the housekeeping gene for normalisation. Expression values were normalised to the corresponding scrambled gRNA control. Statistical significance between enhancer-targeting and scrambled control samples was assessed using an unpaired two-tailed *t*-test with Welch’s correction in GraphPad Prism version 11.0.0. Data are presented as mean ± SEM, and *P* < 0.05 was considered statistically significant. Primer pairs targeting *PAPPA*, *SPRY4*, and *SLC44A2* were obtained from PrimerBank and have been previously validated for quantitative gene expression studies. *ACTB* was used as the housekeeping gene for normalisation of target gene expression. Primer sequences for all targets are as follows:

ACTB_F: CATGTACGTTGCTATCCAGGC
ACTB_R: CTCCTTAATGTCACGCACGAT
PAPPA_F: ACAAAGACCCACGCTACTTTTT
PAPPA_R: CATGAACTGCCCATCATAGGTG
SPRY4_F: TCTGACCAACGGCTCTTAGAC
SPRY4_R: GTGCCATAGTTGACCAGAGTC
SLC44A2_F: AGGCATCATAGCCTGGACTCA
SLC44A2_R: TGGGGAGTGGGACATTGGAA

### RNA-seq processing and quantification

Raw RNA-seq reads were assessed using FastQC v0.12.1^56^ and summarised with MultiQC v1.32^57^. Adapter sequences and low-quality bases were removed using fastp v0.23.4 with automatic paired-end adapter detection enabled. To minimise sequence composition bias at read starts, the first 10 bases of each read were trimmed. Quality assessment was repeated following trimming using FastQC. Trimmed reads were aligned to the human reference genome (GRCh38) using STAR v2.7.10b^58^. Alignments were sorted by genomic coordinate and indexed using SAMtools v1.18^50^. Transcript abundance was quantified independently from the trimmed FASTQ files using Salmon v1.10.2^59^ against a decoy-aware GENCODE v29 transcriptome reference based on GRCh38. Transcript-level abundance estimates were imported into R 4.3.3 using tximeta (v1.20.3)^60^ and aggregated to gene-level counts for downstream analyses.

### Differential gene expression analysis

Differential gene expression analysis was performed in R using DESeq2^61^. Gene-level count matrices generated through tximeta were used as input. Lowly expressed genes were filtered by retaining genes with a mean count ≥ 5 across all samples prior to analysis. Counts were normalised using the median-of-ratios method implemented in DESeq2, and dispersion estimates were calculated using the standard DESeq2 workflow. Differential expression was assessed using a negative binomial generalised linear model with Wald tests, comparing enhancer-targeting CRISPRa samples to their corresponding non-targeting control samples. P-values were adjusted for multiple testing using the Benjamini–Hochberg method, and genes with an adjusted P-value < 0.05 were considered significantly differentially expressed.

### Gene Set Enrichment Analysis

Gene set enrichment analysis (GSEA) was performed using the fgsea package in R (v 1.28.0). Following differential expression analysis, all genes tested by DESeq2 were ranked according to the Wald statistic and used as input for pre-ranked enrichment analysis. Hallmark gene sets obtained from the Molecular Signatures Database (MSigDB)^29^ were used to identify biological pathways associated with enhancer perturbation. Normalised Enrichment Scores (NES) were calculated for each pathway, and statistical significance was determined using Benjamini–Hochberg adjusted *P* values. Pathways with an adjusted *P* value < 0.05 were considered significantly enriched.

### Comparison with publicly available OA cartilage datasets

To assess the relevance of transcriptional changes induced by enhancer perturbation to OA pathology, CRISPRa-derived gene expression signatures were compared with publicly available RNA-sequencing data generated from paired damaged and preserved OA cartilage samples^30^.

Differential expression analysis and GSEA were performed independently on the OA cartilage dataset using the same analytical framework described above. Overlapping differentially expressed genes and shared enriched pathways were subsequently identified to assess concordance between enhancer perturbation and disease-associated transcriptional programmes. Heatmaps were generated using the pheatmap package in R (v 1.0.13), and pathway-level comparisons were visualised using normalised enrichment scores (NES).

## Acknowledgements

The authors would like to acknowledge the assistance given by IT Services and the use of the Computational Shared Facility at The University of Manchester, the study coordinators and the research nurses, and the patients who contributed their time and donated the samples.

## Funding

The portion of the research completed in the United Kingdom is co-funded by the National Institute for Health and Care Research (NIHR) Manchester Biomedical Research Centre (BRC) (NIHR203308), Arthritis UK (award reference 21754), the Medical Research Council (grant number MR/Y030419/1), The views expressed are those of the authors and not necessarily those of the NIHR or the Department of Health and Social Care.

The portion of the research completed in the United States of America was funded by the National Institute of Arthritis and Musculoskeletal and Skin Diseases of the National Institutes of Health under award R21 AR084104. The UNC Flow Cytometry Core Facility (RRID:SCR_019170) is supported in part by the P30 CA016086 Cancer Center Core Support Grant to the UNC Lineberger Comprehensive Cancer Center. The content is solely the responsibility of the authors and does not necessarily represent the official view of the National Institutes of Health. No funding from the NIH was used to support research performed in the United Kingdom.

## Supplementary Methods

### Control oligo selection for MPRA

‘Active control’ sequences were active in MPRA experiments for HEPG2, SK-N-H, and K562 cells and also overlap an ATAC peak in primary human chondrocytes using findOverlaps() from the GenomicRanges R package. Variants with the highest median log2FC across the 3 cell types were included in the library (n=85). We also included tiled oligos across known promoters (CMV, EF1) and genes with biological relevance in the context of OA (ACAN1 and COL2A1) (n=15).

For ‘emVar control’ sequences, selected variants were categorized as emVars in HEPG2, SK-N-H, and K562 cells and also overlap an ATAC peak in chondrocytes (n=50). Since the emVar controls are paired oligos, we generated 2 alleles x 50 emVars for 100 total emVar control sequences in the library. Scrambled versions of the active control sequences were included as negative controls to provide a baseline for low expression levels (n=100). We excluded any variants that contain AsiSI restriction sites or fall within the ENCODE blacklist^62^.

### MPRA Library Construction

All MPRA library construction, barcode–oligo association, RNA capture, and tag-seq library preparation were performed following the Tewhey Lab MPRA v3 protocol (v06/2021, The Jackson Laboratory), with minor modifications described below.

https://www.tewheylab.org/static/pdf/ENCODE_MPRA_Protocol-Tewhey_Lab.pdf

### Oligo Synthesis

Pooled single-stranded DNA oligonucleotides containing candidate regulatory sequences flanked by universal primer binding sites (ACTGGCCGCTTGACG(200bp sequence)CACTGCGGCTCCTGC) were synthesized as an oligo pool (Twist Bioscience). Oligo pools were resuspended in 50 µL nuclease-free water and PCR-amplified using NEBNext® High-Fidelity 2X PCR Master Mix (New England Biolabs) with primers #82_MPRA_v3_F:GCCAGAACATTTCTCTGGCCTAACTGGCCGCTTGACG and #202_MPRA_v3_20I_R:CGACTAGCTTGGCCGCCGACGCTCTTCCGATCT(N1:25252525)(N1) (N1)(N1)(N1)(N1)(N1)(N1)(N1)(N1)(N1)(N1)(N1)(N1)(N1)(N1)(N1)(N1)(N1)(N1)TCTAGAGGTT CGTCGACGCGATCGCAGGAGCCGCAGTG. A test PCR was first performed, and products were visualized on a 1.2% agarose FlashGel to confirm a single discrete band. Full-scale amplification was then performed in twelve parallel 50 µL reactions using 0.5 µL of oligo library per reaction. Cycling conditions were 98°C for 30 s; 6 cycles of 98°C for 10 s, 60°C for 15 s, and 65°C for 45 s; followed by 72°C for 5 min. PCR products were purified using 0.8× CleanNGS DNA C RNA Clean-Up Magnetic Beads (Bulldog Bio) and eluted in nuclease-free water. Library size (∼329 bp) was verified on a 2.2% agarose FlashGel, and DNA concentration was measured by NanoDrop spectrophotometry.

### Vector Assembly

pMPRAv3:Δluc:ΔxbaI was a gift from Pardis Sabeti C Ryan Tewhey (Addgene plasmid # 109035, http://n2t.net/addgene:109035; RRID:Addgene_109035). For construction of the mpraΔorf library, 10 µg of pMPRAv3Δluc:ΔxbaI was digested with SfiI (New England Biolabs) in CutSmart buffer and incubated at 50°C overnight. Digested backbone was purified using 1.5× CleanNGS DNA C RNA Clean-Up Magnetic Beads (Bulldog Bio) and verified by agarose gel electrophoresis.

Gibson assembly reactions were performed using NEB HiFi DNA Assembly Master Mix (New England Biolabs) with 2.2 µg purified oligo insert and 2 µg digested backbone in a 100 µL reaction incubated at 50°C for 1 hour. Assemblies were purified using 1.2× CleanNGS beads and eluted in 20 µL Lo-Bind tubes.

The assembly product (10ng) was electroporated into 50 µL 10-beta electrocompetent E. coli (New England Biolabs) using 2 kV, 200 Ω, 25 µF settings, and immediately recovered in 4 mL SOC for 1 hour at 37°C with agitation (200 r.p.m.). A 2 µL aliquot was diluted in 400 µL LB and spread on a prewarmed 15-cm LB-ampicillin agar plate to determine colony-forming units (CFU). The remaining culture was split between eight prewarmed 15-cm LB-ampicillin agar plates and incubated overnight at 37°C. After CFU counting, two plates were scraped with 6 mL LB and combined for purification using the ZymoPURE II Plasmid Maxiprep Kit (Zymo Research). Library insertion was confirmed by colony PCR and Sanger sequencing of representative clones.

### Barcode–Oligo Association

Plasmid library DNA (200 ng) was PCR-amplified using primers #197_MPRA_v3_TruSeq_Amp2Sa_F:GTGACTGGAGTTCAGACGTGTGCTCTTCCGATCTAACT GGCCGCTTGACG and #757_Ilmn_P5_1stPCR: ACACTCTTTCCCTACACGACGCTCTTCCGATC with NEBNext® High-Fidelity 2X PCR Master Mix. Cycling conditions were 98°C for 30 s; 5 cycles of 98°C for 20 s, 62°C for 15 s, and 72°C for 30 s; followed by 72°C for 2 min. Amplicons were purified using 1× CleanNGS beads and eluted in 30 µL EB.

Indexing adapters were added by a second PCR using P5 and P7 index primers with cycling conditions of 98°C for 30 s; 5 cycles of 98°C for 10 s, 62°C for 15 s, and 72°C for 30 s; followed by 72°C for 2 min. Indexed libraries were purified using 1× CleanNGS beads, quantified, pooled by molarity, and sequenced on an Illumina platform using paired-end 150 bp reads.

### Barcode mapping

MPRAsnakeflow v0.4.2 (https://github.com/kircherlab/MPRAsnakeflow/tree/v0.4.2) was used to pair barcodes with each oligo from the MPRA library^26^. Required inputs include paired-end DNA sequencing (.fastq) and designed oligos from the MPRA library (.fasta) for mapping. Barcode-oligo pairs were mapped using an exact-match approach, such that oligos had to match the designed sequence at every position across 200bp. MPRAsnakeflow config.yaml specified bc_length: 20, tool: exact, sequence_length: 200, alignment_start: 1, and default for other parameters. The resulting assignment_barcodes.default.tsv.gz file was used as the assignment file in the MPRAsnakeflow experiment workflow below.

### Generation of mpra:gfp Library

pMPRAv3:minP-GFP was a gift from Pardis Sabeti C Ryan Tewhey (Addgene plasmid # 109036; http://n2t.net/addgene:109036; RRID:Addgene_109036)

mpraΔorf plasmid was linearized with AsiSI (New England Biolabs) at 37°C overnight and purified using Monarch DNA Cleanup Columns (New England Biolabs).

A GFP amplicon was generated by PCR from pMPRAv3:minP-GFP (pMPRAv3:minP-GFP, Addgene plasmid #109036) using primers #200_MPRA_v3_GFP_Fusion_v2_F: CACTGCGGCTCCTGCGATCTAACTGGCCGGTACCTGAGCTCGCTA and #201_MPRA_v3_GFP_Fusion_v2_R:TCTAGAGGTTCGTCGACGCGATTATTATCATTACTTGTAC AGCTCGTCCATGC with NEBNext® High-Fidelity 2X PCR Master Mix. Cycling conditions were 98°C for 30 s; 20 cycles of 98°C for 10 s, 60°C for 15 s, and 72°C for 45 s; followed by 72°C for 5 min. PCR products were treated with DpnI at 37°C for 30 min and purified by double-SPRI cleanup using CleanNGS beads (0.5× then 1.5×), eluting in 40 µL.

A second amplification (GFP:amp2) was performed using the same primers and cycling conditions. Reactions were pooled and purified using 1.5× CleanNGS beads.

GFP amplicons were inserted into the linearized mpraΔorf library by Gibson assembly using NEB HiFi DNA Assembly Master Mix and incubated at 50°C for 90 min. Assemblies were purified using 1.5× CleanNGS beads. Remaining uncut vector was removed by digestion with AsiSI and RecBCD (New England Biolabs) at 37°C overnight, followed by 1.5× CleanNGS purification.

The assembly product (10 ng) was electroporated into 50 µL 10-beta electrocompetent *E. coli* (New England Biolabs) using 2 kV, 200 Ω, 25 µF settings, and immediately recovered in 4 mL SOC for 1 hour at 37°C with agitation (200 r.p.m.). A 2 µL aliquot was diluted in 400 µL LB and spread on a prewarmed 15-cm LB-ampicillin agar plate to determine colony-forming units (CFU). The remaining culture was split between eight prewarmed 15-cm LB-ampicillin agar plates and incubated overnight at 37°C. After CFU counting, two plates were scraped with 6 mL LB and combined for purification using the ZymoPURE II Plasmid Maxiprep Kit (Zymo Research). Library insertion was confirmed by colony PCR and Sanger sequencing of representative clones.

### RNA Extraction G cDNA Synthesis

Cells transfected with mpra:gfp libraries were harvested and total RNA was extracted using the RNeasy Mini Kit (Qiagen) according to the manufacturer’s instructions. RNA was treated with TURBO DNase Free Kit (Thermo Fisher Scientific).

GFP transcripts were captured by hybridization with a GFP 2-probe biotinylated probe mix (10 µM total) in 20× SSC and deionized formamide at 65°C for 2.5 hours. Probes were GFP_BiotinCapture1:CCAGGATGTTGCCGTCCTCCTTGAAGTCGATGCCC/3BioTEG/ and GFP_BiotinCapture2:CCTCGATGTTGTGGCGGGTCTTGAAGTTCACCTTG/3BioTEG/. Hybridized RNA was captured using Dynabeads MyOne Streptavidin C1 beads (Thermo Fisher Scientific), washed once with 1× SSC and twice with 0.1× SSC, and eluted in nuclease-free water containing SUPERase In (Thermo Fisher Scientific). Captured RNA underwent an additional TURBO DNase treatment and was purified using 2× CleanNGS beads.

First-strand cDNA synthesis was performed using SuperScript III First-Strand Synthesis SuperMix kit (Thermo Fisher Scientific) with gene-specific primer #19 (CCGACTAGCTTGGCCGC). The primer annealing was carried out at 65°C for 5 minutes and synthesis was carried out at 47°C for 80 min followed by enzyme inactivation at 85°C. The cDNA was purified using 2× CleanNGS beads and eluted in EB.

### Tag-seq Library Preparation for Illumina Sequencing

To determine optimal PCR cycle number, qPCR was performed on serial dilutions of plasmid library (standard curve) and GFP cDNA using NEBNext® High-Fidelity 2X PCR Master Mix with primers

MPRA_v3_Illumina_GFP_F:GTGACTGGAGTTCAGACGTGTGCTCTTCCGATCTCGCCCTGAG CAAAGACC and

TruSeq_Universal_Adapter:AATGATACGGCGACCACCGAGATCTACACTCTTTCCCTACACG ACGCTCTTCCGATCT and SYBR Green I (Thermo Fisher Scientific). Cycling conditions used were 98C for 30s; 40x {98C for 10s; 62C for 15s; 72C for 30s}; 72C for 2m; Melt Curve Analysis.

First-round PCR amplification of barcode regions was performed using NEBNext® High-Fidelity 2X PCR Master Mix with cycle numbers determined from qPCR. Products were purified using 1× CleanNGS beads and eluted in EB. A second PCR was performed to add Illumina P5 and P7 index primers. Indexed libraries were purified using 1× CleanNGS beads, quantified by Qubit fluorometry (Thermo Fisher Scientific), assessed by TapeStation (Agilent), pooled by molarity, and sequenced on an Illumina platform using single-end 26 bp reads.

### MPRA Analysis

Barcode-level RNA and DNA count matrices were generated using the MPRAsnakeflow experiment workflow v0.4.2 with barcode–oligo assignment files from the mapping workflow. Downstream analyses were performed using a custom R pipeline available on github (https://github.com/srstuppy/MPRA).

Barcode-level RNA and DNA counts were filtered to remove low-RNA observations and jackpot barcodes (barcodes contributing >50% of the total RNA counts for a given oligo within a donor × condition). Barcode activity was calculated as the log2 ratio of RNA to DNA counts and median-centred within each donor × condition using scrambled negative-control oligos. Barcode-level activity values were aggregated to the oligo level using medians.

Allelic effects were assessed by pairing reference and alternate oligos for each rsID and comparing activity distributions between alleles using two-sided t-tests. One-pass Grubbs outlier removal was applied separately within each allele prior to testing. Variants with at least two observations per allele were tested.

Multiple testing correction was performed using the Benjamini–Hochberg (BH) procedure (FDR < 0.1). BH adjustment was applied globally across all variant pairs, among variants with sufficient RNA support (total RNA across ref+alt >24), among emVar control pairs only, and within loci. Variants below the RNA-support threshold were retained in result tables but excluded from BH testing. Analyses were performed separately for PBS, Fn-f, and combined datasets.

### Reporter Assays

Individual test sequences were synthesized as double-stranded DNA gene fragments (Twist Bioscience) containing Gibson homology arms matching the pMPRAv3:minP-GFP backbone. The pMPRAv3:minP-GFP vector (Addgene #109036) was linearized with BglI in 10X NEBuffer r3.1 (New England Biolabs) and purified using Zymo DNA Clean C Concentrator columns (Zymo Research). Inserts were assembled into the linearized backbone using NEB HiFi DNA Assembly Master Mix (New England Biolabs). Assembly reactions were chemically transformed into NEB Stable competent E. coli (New England Biolabs), and plasmid DNA was isolated using Zymo miniprep kits. Correct constructs were confirmed by whole-plasmid sequencing (Plasmidsaurus). Sequence-verified plasmids were expanded and purified using Zymo maxiprep kits for downstream nucleofection.

Low-throughput reporter assays were performed in primary human chondrocytes using the same isolation, nucleofection, and treatment conditions described for MPRA transfections. Reporter activity was assessed by flow cytometry and immuno fluorescence (IF) using both native and antibody-based detection of GFP. For both assays, cells were fixed with 4% PFA, permeabilized with 0.03% Triton X-100, blocked with BSA, and stained with a directly conjugated anti-GFP monoclonal antibody (Invitrogen/eBioscience, clone 5F12.4, eFluor™ 660; Cat. No. 50-6498-82) and counterstained with DAPI to label nuclei. For flow cytometry, stained cells were analysed to quantify the percentage of GFP-positive cells and/or GFP fluorescence intensity using standard gating strategies, with DAPI used to gate nucleated events. For IF, stained cells were imaged under identical acquisition settings across conditions, and GFP fluorescence was quantified per cell using CellProfiler.

---

**Supplementary Figure 1.**
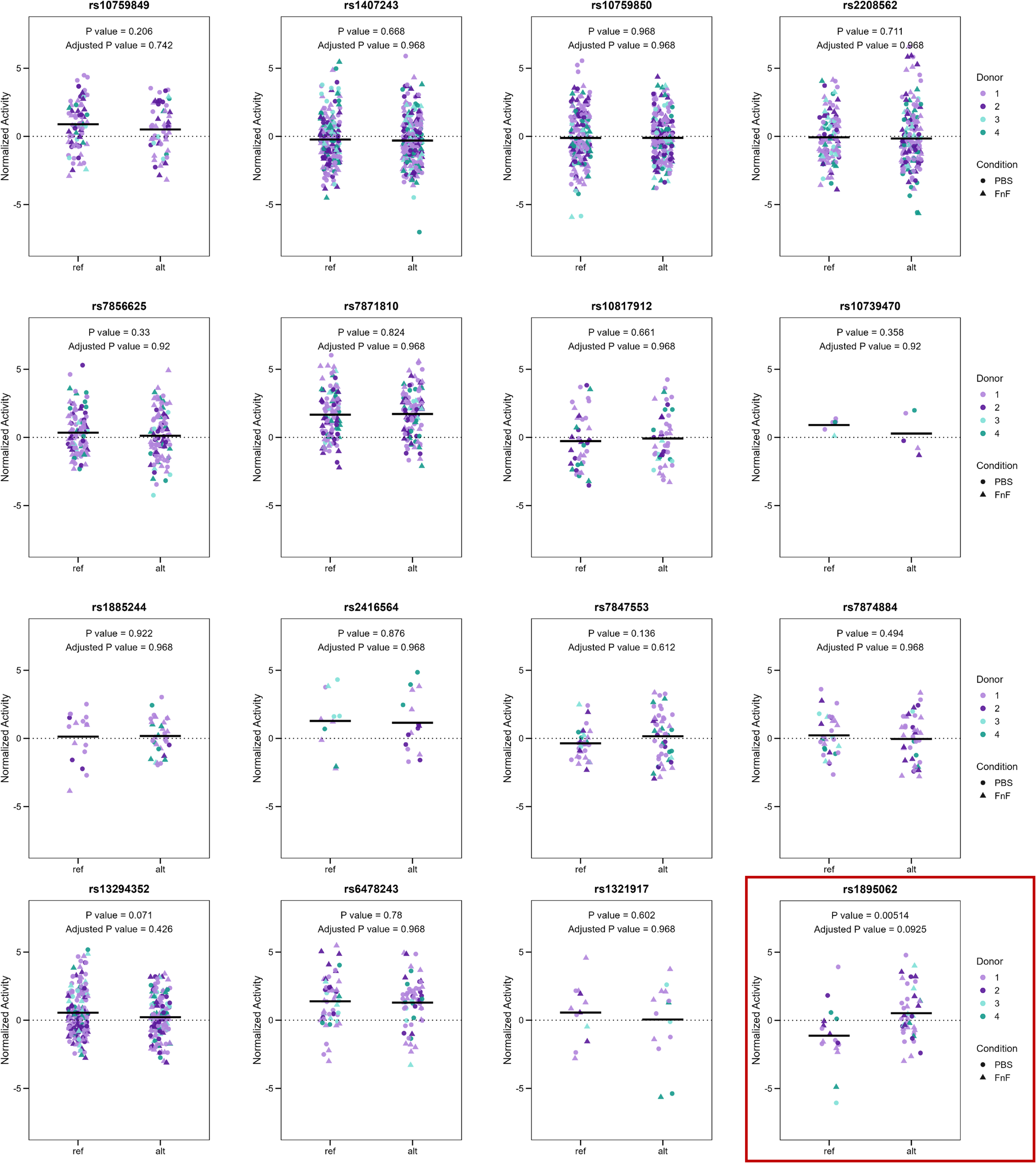
MPRA activity of variants at the *PAPPA* locus. Normalized reporter activity for the reference (ref) and alternative (alt) alleles of 15 additional variants at the *PAPPA* locus. Individual points represent barcode-level measurements, colored by donor (1–4) and shaped by treatment condition (PBS, circles; Fn-f, triangles). Horizontal black lines indicate median normalized activity. P values were calculated using two-sided t-test comparing reference and alternative allele activity, with Benjamini–Hochberg correction for multiple testing. Nominal and adjusted P values are displayed for each variant, with an adjusted P value < 0.10 considered statistically significant. Bottom right in red box shows rs1895062 (reproduced from Fig. 3A).

**Supplementary Figure 2.**
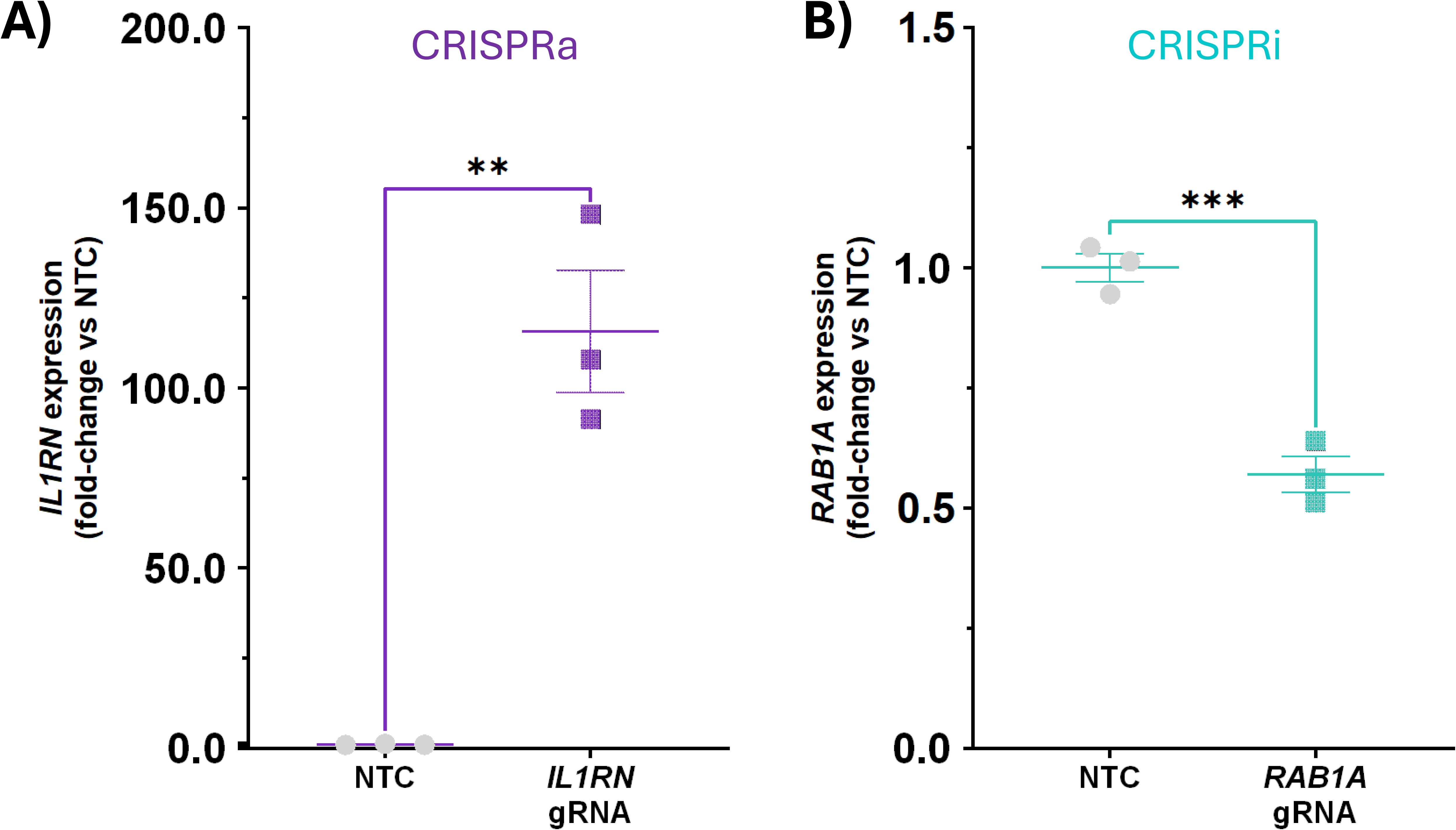
Functional validation of stable CRISPRa and CRISPRi SW1353 cell lines. **(A)** Validation of CRISPRa-mediated transcriptional activation following transduction of an *IL1RN*-targeting guide RNA into SW1353-CRISPRa cells. *IL1RN* expression was quantified by qPCR and normalised to *ACTB* and the corresponding scrambled guide RNA control. Targeting of the *IL1RN* promoter resulted in a 115-fold increase in *IL1RN* expression relative to scrambled controls (*P* = 0.0210). **(B)** Validation of CRISPRi-mediated transcriptional repression following transduction of a *RAB1A*-targeting guide RNA into SW1353-CRISPRi cells. *RAB1A* expression was quantified by qPCR and normalised to *ACTB* and the corresponding scrambled guide RNA control. Targeting of the *RAB1A* promoter resulted in an approximately 45% reduction in *RAB1A* expression relative to scrambled controls (*P* = 0.0011). Data are presented as mean ± SEM from three independent biological replicates. Statistical significance was assessed using an unpaired two-tailed *t*-test with Welch’s correction. ***P* < 0.05 (*), *P* < 0.01 (**).**

**Supplementary Fig 3.**
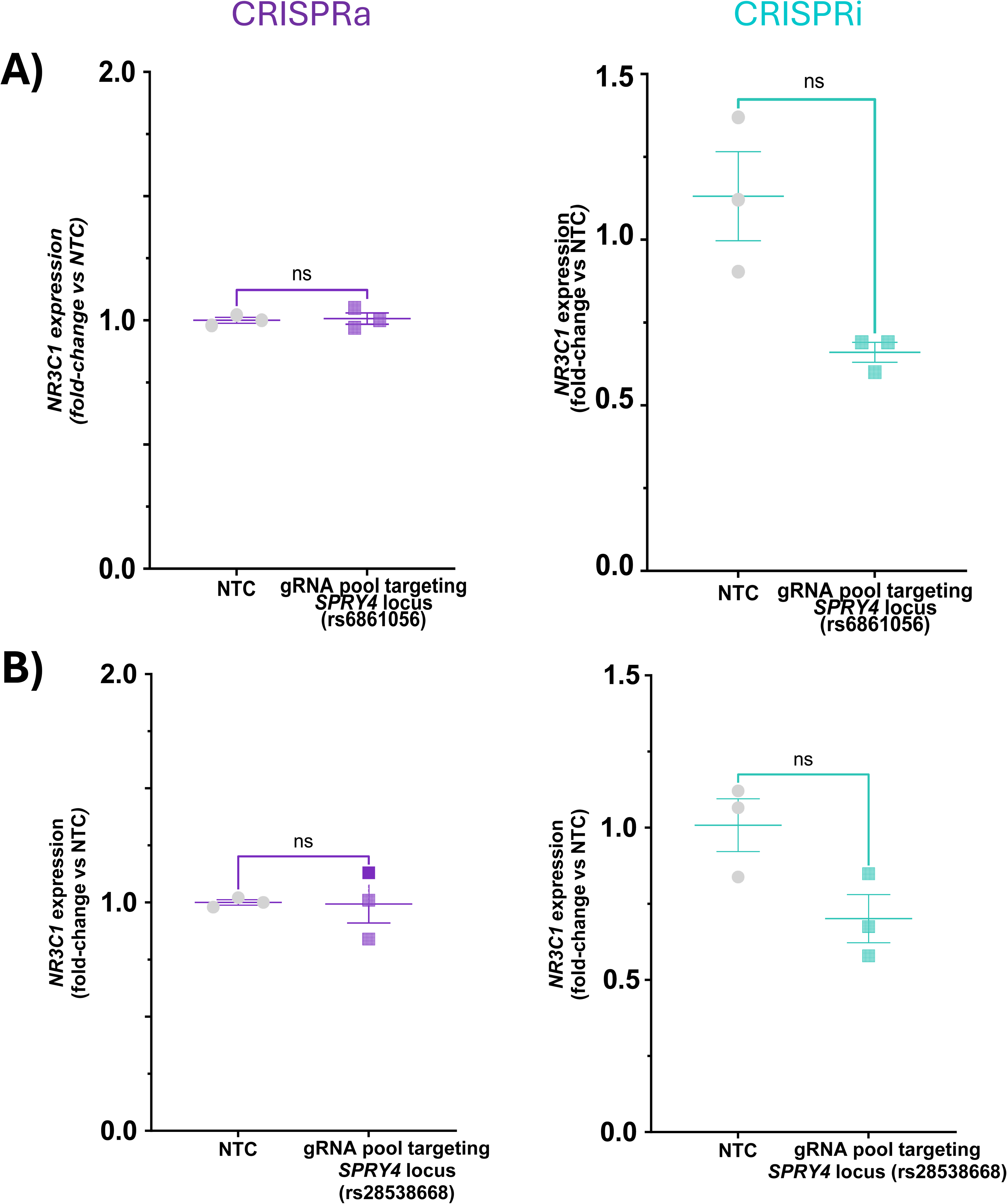
Effect of activating and inhibiting SPRY4 locus variants on NR3C1 expression. **(A)** Quantitative PCR analysis of NR3C1 expression following CRISPRa (left panel) and CRISPRi (right panel) perturbations targeting the OA-associated variant rs6861056 at the SPRY4 locus. Expression levels are shown relative to NTC. No statistically significant changes in NR3C1 expression were detected. **(B)** Quantitative PCR analysis of NR3C1 expression following CRISPRa (left panel) and CRISPRi (right panel) targeting of a second candidate regulatory element at the SPRY4 locus containing variant rs28538668. Expression levels are shown relative to NTC. No statistically significant changes in NR3C1 expression were observed. Individual points represent biological replicates and horizontal bars indicate mean ± SEM. Statistical significance was determined by comparison with NTC controls. P < 0.05; ns, not significant.

**Supplementary Figure 4.**
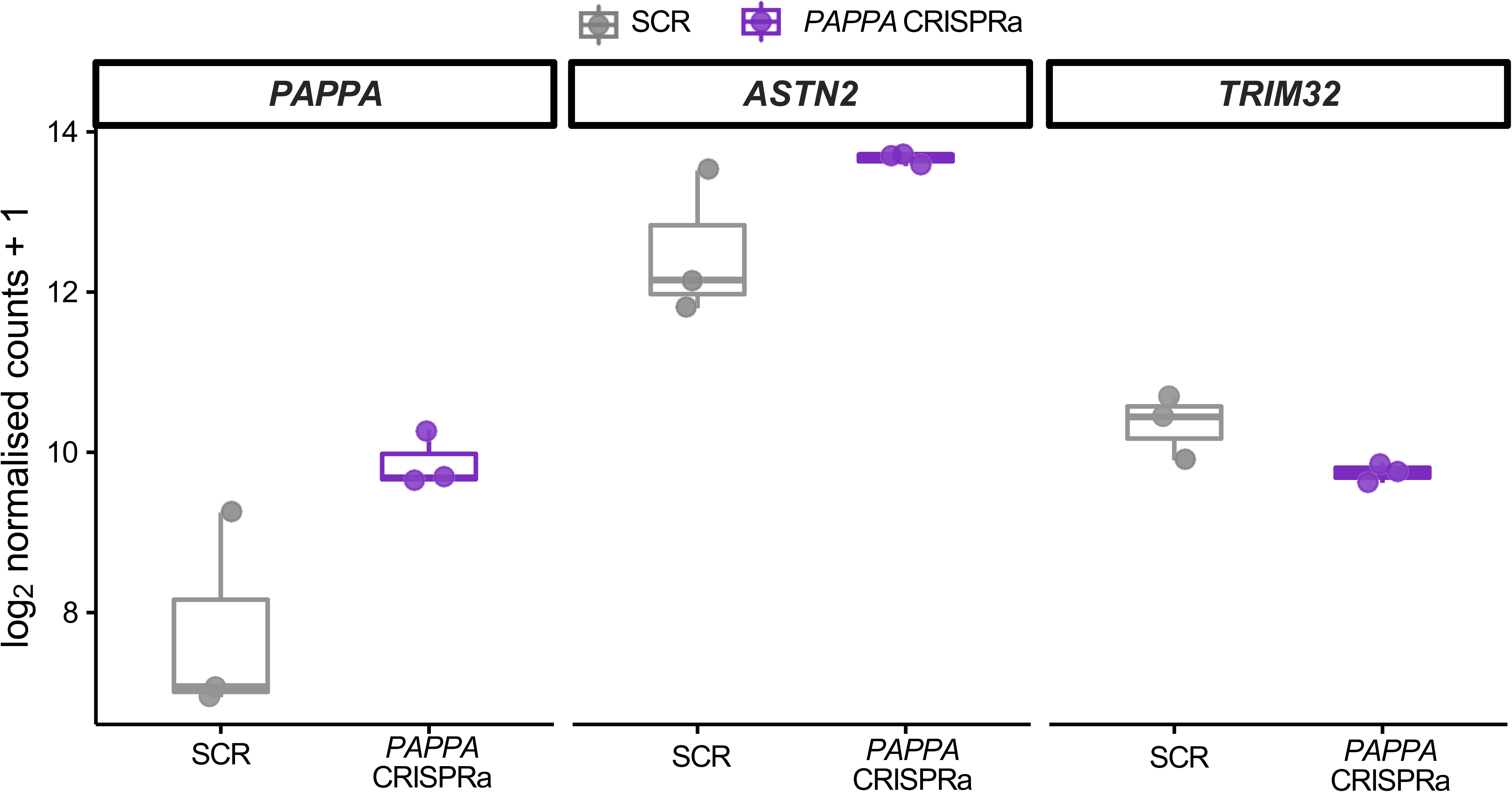
Expression of PAPPA and neighboring genes following CRISPRa targeting the rs18G5062-containing regulatory region. Expression of PAPPA, ASTN2, and TRIM32 following CRISPR activation (CRISPRa) of the regulatory region containing rs1895062, compared with a scrambled guide RNA control (SCR). Gene expression is presented as log2-normalized counts + 1. Individual points represent replicate measurements, with boxplots indicating the median and interquartile range.

## Notes

### Competing Interest Statement

The authors have declared no competing interest.

